# LGL-1 and the RhoGAP protein PAC-1 redundantly polarize the *C. elegans* embryonic epidermis

**DOI:** 10.1101/2025.09.05.674236

**Authors:** Olga D. Jarosińska, Amalia Riga, Hala Zahreddine Fahs, Joren M. Woeltjes, Ruben Schmidt, Fathima S. Refai, Suma Gopinadhan, Kristin C. Gunsalus, Mike Boxem

## Abstract

Apical–basal polarity is essential for epithelial organization and function and is established by conserved cortical polarity proteins. However, the requirements for canonical polarity factors vary between tissues and organisms. For example, the basolateral protein lethal giant larvae (Lgl) is essential for epithelial polarity in *Drosophila* but dispensable in *C. elegans*. To better define the epithelial polarity program in *C. elegans*, we performed a genome-wide RNAi screen for synthetic lethality with *lgl-1*. Combined loss of LGL-1 and the RhoGAP PAC-1 caused embryonic lethality due to elongation defects and epidermal rupture. Epidermal cells showed expansion of the apical domain and aPKC, accompanied by mislocalization of junctional proteins and LET-413^Scribble^. These defects indicate overactivity of apical polarity determinants. Consistently, partial inactivation of aPKC or CDC-42 suppressed lethality in *pac-1; lgl-1* animals. Together, our results identify PAC-1 and LGL-1 as redundant inhibitors of apical polarity in the *C. elegans* embryonic epidermis and provide new insights into how conserved mechanisms are adapted across species.

## Introduction

The form and functioning of epithelial cells are dependent on their organization along an apical–basal axis of polarity. Epithelial polarity is established by conserved polarity regulators that engage in mutually antagonistic interactions to establish distinct apical, lateral, and basal membrane domains, and position celjunctions (1–4). Specification of the apical domain involves the activities of the partitioning defective proteins Par3 (Bazooka in *Drosophila*) and Par6, atypical protein kinase C (aPKC), the small Rho GTPase Cdc42, the transmembrane protein Crumbs, and its intracellular adapter Pals1 (Stardust in *Drosophila*). Formation of the basolateral domain depends on the Scribble module proteins Scribble (Scrib), Discs large (Dlg), and Lethal giant larvae (Lgl), as well as on the kinase Par1.

Mutual inhibition between aPKC and Lgl plays a central role in the network of antagonistic interactions that establish apical–basal polarity. At the apical domain, aPKC phosphorylates a range of polarity proteins to mediate their exclusion from the plasma membrane, including Lgl (5–9). aPKC strongly interacts with Par6, and formation of a Par6/aPKC complex is necessary for both localization and activity of aPKC (10–12). Par6 can locate aPKC to the apical membrane domain by interacting with Par3 or Cdc42, both of which can bind Par6 and recruit Par6/aPKC to the membrane (11,13–18). Full activation of the kinase activity of aPKC is promoted by binding to Cdc42, and in the *C. elegans* zygote Par6/aPKC have been shown to shuttle between an inactive PAR-3 complex and an active CDC-42 complex [11,12,14,19–22]. Binding of Par6/aPKC to Cdc42 also increases the affinity of Par6 for Crumbs, and handoff from Cdc42 to Crumbs may further stabilize active Par6/aPKC at the membrane (15,20,23). At the basolateral domain, Lgl is required to antagonize the activity of Par6/aPKC (24–26). Lgl forms a ternary complex with Par6/aPKC via multiple interaction interfaces (8,27,28). However, it is still unclear if the antagonistic activity towards aPKC is based on inhibition of kinase activity or if Lgl prevents phosphorylation of other aPKC targets by competitively keeping aPKC engaged with processive Lgl phosphorylation.

While these cortical polarity determinants are highly conserved in metazoans, much of our understanding of their functioning in apical–basal polarity establishment in epithelia is based on studies in *Drosophila*. It is clear, however, that there is considerable variation in the mechanisms that establish apical–basal polarity between different organisms and between different epithelial tissues in the same organism. For example, in *Drosophila*, not all epithelial tissues that express Crumbs depend on it to establish polarity (29–31), while all three *C. elegans* Crumbs family members are collectively dispensable for polarity establishment (32,33). Recent studies in *Drosophila* have also shown that the enterocytes of the midgut do not require any of the canonical polarity factors and appear to use a different mode of polarization (34). It is important therefore to investigate the mechanisms that underly apical–basal polarization in different developmental and organismal contexts, and determine how universal the roles of different polarity modules are.

The *C. elegans* Lgl ortholog LGL-1 is a striking example of how polarity proteins may vary in their importance in epithelial polarization. While *Drosophila* Lgl is essential for epithelial polarity establishment, *C. elegans* lacking LGL-1 are viable and show no defects in polarity establishment. (35,36). In the one cell embryo, the lack of an essential role for LGL-1 may be explained by the presence of the *C. elegans* specific protein PAR-2, which acts redundantly with LGL-1 (35,36). However, expression of PAR-2 is limited to the germ lineage while LGL-1 is broadly expressed (35–37). LGL-1 may also antagonize aPKC in the spermatheca, as *lgl-1* mutants were found as suppressors of a temperature sensitive *aPKC* allele that causes junction breaks in the spermatheca and leads to sterility (38). It is not known, however, whether LGL-1 functions similarly in other epithelia or why loss of LGL-1 does not cause epithelial polarity defects.

Another polarity regulator whose functioning may differ between *Drosophila* and *C. elegans* is the RhoGAP protein PAC-1 (RhoGAP19D in *Drosophila* and ARHGAP21 in mammals). PAC-1 was identified as a regulator of radial polarization of *C. elegans* blastomeres, mediating the asymmetric enrichment of Par6/aPKC to the outer contact-free cell surfaces (39). E-cadherin-dependent recruitment of PAC-1 to sites of cell–cell contact is thought to locally inactivate CDC-42, restricting recruitment of Par6/aPKC by CDC-42 to the outer surfaces (39,40). Additionally, in epidermal cells of the embryo, PAC-1 is involved in regulating junctional actin organization and the levels of AJ proteins, likely also by regulating CDC-42 activity (41). However, *pac-1* is not an essential gene. Loss of *pac-1* causes defects in cell positioning during gastrulation but >90% of embryos hatch and develop normally (41). In contrast, *Drosophila* RhoGAP19D was identified as an essential protein that antagonizes aPKC at the lateral domain (42). RhoGAP19D is recruited to the lateral domain of follicle cells by the E-cadherin complex, and its loss leads to ectopic lateral Cdc42 activity, increased Par6/aPKC activity, and expansion of the apical domain. *rhogap19d* mutants die at various stages of development, indicating a broad role of RhoGAP19D in epithelial polarization (42).

Here, we investigate the role of LGL-1 in polarity establishment in *C. elegans*. We performed an RNAi-based synthetic lethality screen in an *lgl-1* deletion background and found that the combined loss of *lgl-1* and *pac-1* leads to a fully penetrant lethal phenotype, with most animals arresting at the 2-fold stage of embryonic elongation with frequent rupturing of the epidermis. The rupturing is likely due to severe defects apical–basal polarization and apical junction formation in the epidermis. We show that the lethality of combined *lgl-1* and *pac-1* loss can be reduced by lowering the levels or activity of the apical polarity proteins CDC-42 or aPKC. These experiments are the first demonstration that LGL-1 and PAC-1 act together in epithelial polarization in *C. elegans*, likely by redundantly controlling aPKC activity. As LGL-1 has now been shown to act redundantly with a second polarity regulator in two tissues, we speculate that such redundancy is a general feature of LGL-1 in *C. elegans*. Our data thus imply that differences in essentiality of cortical polarity regulators reflect changes in the strengths of the wiring of the different connections in the epithelial polarity network, rather than major differences in the mode of polarization.

## Results

### The RhoGAP protein PAC-1 acts redundantly with LGL-1 in embryonic development

Previously, LGL-1 was shown to act redundantly with PAR-2 during polarity establishment in the one cell embryo. To identify genes that may act redundantly with *lgl-1* in other developmental stages, we performed an RNAi-based enhancer screen. We first generated the *lgl-1(mib44)* deletion allele by replacing most of the *lgl-1* coding sequence with *GFP* expressed from the *myo-2* promoter (Fig. S1A). We then used this strain to perform a whole-genome feeding RNAi screen using the Ahringer feeding library in an automated robotics pipeline (43,44) (Fig. 1A). We identified 79 RNAi clones that caused stronger fitness or developmental defects in *lgl-1(mib44)* animals than in wild-type controls. We discarded 27 clones frequently identified as enhancers in the screening pipeline as likely false positives. Of the remaining clones, 5 retested positive by feeding RNAi on plates using sequence-verified RNAi clones (Fig. S1B). We decided to focus on the RhoGAP encoding gene *pac-1* because of the strong synthetic lethality displayed with *lgl-1* and its previously described roles in radial polarization of early embryonic blastomeres and regulation of CDC-42 [28,39,45,45].

**Figure 1.**
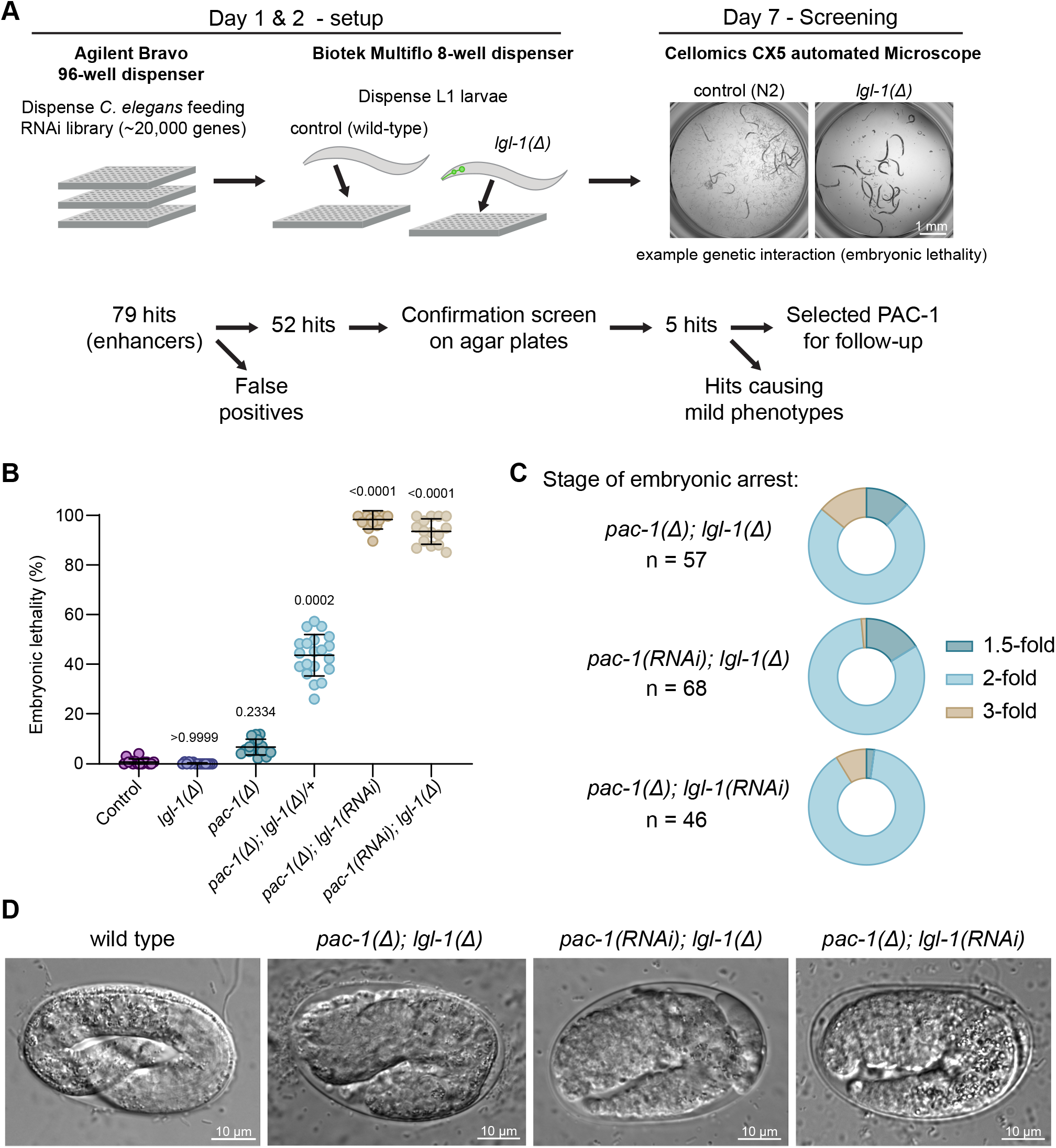
The RhoGAP protein PAC-1 acts redundantly with LGL-1 in embryonic development. (**A**) Overview of the genome-wide RNAi screen procedure. (**B**) Embryonic lethality observed in progeny of animals of indicated genotypes. Data is represented as mean ± SD and analyzed with a Kruskal-Wallis test followed by Dunn’s multiple pairwise comparison test; p values shown on graph. Total plates counted in order of samples in graph: 15, 15, 15, 20, 12 and 15. Total progeny counted in order of samples in graph: 1683, 1726, 1670, 1944, 928, and 713. (**C**) Stage of embryonic arrest of embryos of indicated genotypes. Homozygous *pac-1(mib269); lgl-1(mib44)* embryos were derived from *pac-1(mib269); lgl-1(mib44)/+* parents. n = number of embryos counted. (**D**) Representative DIC images of wild-type, *pac-1(mib269); lgl-1(mib44), pac-1(RNAi); lgl-1(mib201)*, and *pac-1(mib269); lgl-1(RNAi)* embryos. Strains used: CGC1, BOX1221, BOX919 and BOX907.

To further investigate the interaction between *pac-1* and *lgl-1*, we generated the *pac-1(mib269)* deletion allele, which removes the entire *pac-1* coding sequence. We also generated a second *lgl-1* deletion allele *(mib201)*, replacing the GFP in *mib44* with BFP to mitigate imaging problems due to the extreme brightness of *Pmyo-2::GFP* (Fig. S1A). The *lgl-1* and *pac-1* deletion strains are viable and behave as previously described strains carrying loss-of-function alleles. Deletion of *lgl-1* does not cause apparent developmental defects (35,36), while loss of *pac-1* causes ∼10% embryonic lethality with pleotropic phenotypes, while the remaining animals developing normally (Fig. 1B, S1C) (39,41). In contrast, injection of *pac-1* dsRNA in *lgl-1(mib201)* or of *lgl-1* dsRNA in *pac-1(mib269)* homozygous mutants resulted in nearly complete embryonic lethality (Fig. 1B). To further confirm the genetic interaction, we generated animals homozygous for *pac-1(mib269)* and heterozygous for *lgl-1(mib44)* and examined their offspring. Interestingly, progeny from these *pac-1(mib269); lgl-1(mib44)/+* parents showed 43% embryonic or early larval lethality rather than the expected 25%, indicating that the *lgl-1* deletion allele is partially haplo-insufficient (Fig. 1B). Consistent with this interpretation, 45% of the surviving animals expressed GFP, and hence carried at least one copy of the *lgl-1* deletion allele, rather than the expected 67%.

Differential interference contrast (DIC) microscopy of *pac-1(mib269); lgl-1(mib44)* embryos derived from *pac-1; lgl-1/+* parents revealed that 73% of embryos arrest at the 2-fold stage of embryonic elongation (Fig. 1C), with a bulging epidermis and frequent ruptures of the epidermis (Fig. 1D). To assess whether maternal *lgl-1* contribution might mask an earlier phenotype, we depleted *pac-1* by RNAi in homozygous *lgl-1(mib44)* mutants and depleted *lgl-1* in homozygous *pac-1(mib269)* animals. Both treatments resulted in >80% of embryos arresting at the 2-fold stage (Fig. 1C, D), indicating that combined maternal and zygotic loss of *pac-1* and *lgl-1* results in a predominantly 2-fold arrest. The remaining embryos still arrested, but at slighlty earlier or later timepoints in embryonic development, likely reflecting inherent biological variation. Subsequent experiments were done with RNAi for either *pac-1* or *lgl-1* in the genetic null background for the other gene.

Loss-of-function mutants that strongly enhance a phenotype are often interpreted as acting in parallel pathways. We therefore examined whether loss of *lgl-1* or *pac-1* alters the localization of endogenously GFP-tagged LGL-1 or PAC-1. In neither null background did we detect changes in the subcellular localization of the other protein, consistent with LGL-1 and PAC-1 functioning in parallel pathways (Fig. S1D). Together, these experiments show that *pac-1* and *lgl-1* are redundantly required for embryonic development.

### PAC-1 and LGL-1 redundantly control the positioning of apical junctions

Embryonic elongation is dependent on adherens junctions, and the defects we observe in embryos lacking *lgl-1* and *pac-1*, including the 2-fold arrest and rupturing of the epidermis, are also observed in embryos with defects in junctional integrity (46–48). Moreover, loss of PAC-1 was previously shown to block elongation in embryos carrying the hypomorphic HMP-1/α-catenin mutant *fe4*, which has reduced actin binding capability (41,49). We therefore investigated if combined loss of *pac-1* and *lgl-1* results in defects in junctional integrity in embryonic epidermal cells. We visualized the distribution of each of the three junctional complexes (50) in *pac-1(RNAi); lgl-1(mib201)* embryos using a representative protein endogenously fused to a fluorescent protein. We imaged HMR-1::GFP (E-cadherin) from the cadherin/catenin complex (CCC), AFD-1::GFP (Afadin) from the SAX-7/MAGI-1/AFD-1 complex (SMAC) and DLG-1::mCherry from the DLG-1/AJM-1 complex (DAC) (50).

Consistent with our DIC observations, time-lapse imaging of DLG-1::mCherry or the membrane marker PH::GFP in *pac-1(RNAi); lgl-1(mib201)* embryos showed that double mutant embryos completed ventral closure (10/12 embryos imaged) and arrested during elongation with rupturing of the epidermis visible as expulsion of internal cells (Fig. S2A and Video S1). We observed ruptures on both the ventral and dorsal side of the embryos, and at different positions along the anterior–posterior body axis. Thus, the weakening of the epidermis does not appear to be limited to any particular region.

In 3D confocal imaging, we observed aberrant lateral spreading of each of the three junctional proteins in >90% of embryos examined (visible as double lines in maximum intensity projections, see arrowheads in Fig. 2A, and quantified in Fig. 2B). The defects were most severe for DLG-1, which also showed gaps in the normally continuous junctional band (arrows in Figs. 2A and S2B). Imaging of embryos in earlier stages of development showed that normal continuous junctional DLG-1 bands are never established in *pac-1(RNAi); lgl-1(mib201)* embryos (Fig. S2B). To better visualize the basolateral mislocalization, we imaged DLG-1 together with PH::GFP. In cross-sections of *pac-1(RNAi); lgl-1(mib201)* embryos, DLG-1 was spread along the lateral membrane compared to the tight subapical localization observed in controls expressing wild-type *lgl-1* and *pac-1* (Fig 2C, D). Finally, we rendered image stacks of embryos expressing HMR-1::GFP and DLG-1::mCherry in 3D (Fig. 2E). Control animals showed the expected subapical localization of both proteins, with HMR-1 signal appearing slightly more apical than DLG-1 (Fig. 2F - junctions ‘i’ and ‘ii’, and Video S2). However, in *pac-1(RNAi); lgl-1(mib201)* embryos, we observed severe defects in DLG-1 and HMR-1 localization. First, ∼25% of junctions showed lateral displacement of DLG-1 and HMR-1 along the lateral membrane (Fig. 2F - junction ‘iii’, Fig. 2G, and Video S2). The two proteins did not mix completely but showed remaining organization with regions expressing predominantly DLG-1 being flanked by HMR-1 signal. Second, in ∼50% of junctions we observed ring-like structures ofjunctional material encircling a junction-free area on the lateral domain (Fig. 2F, junction ‘iv’, and Fig. 2G).

**Figure 2.**
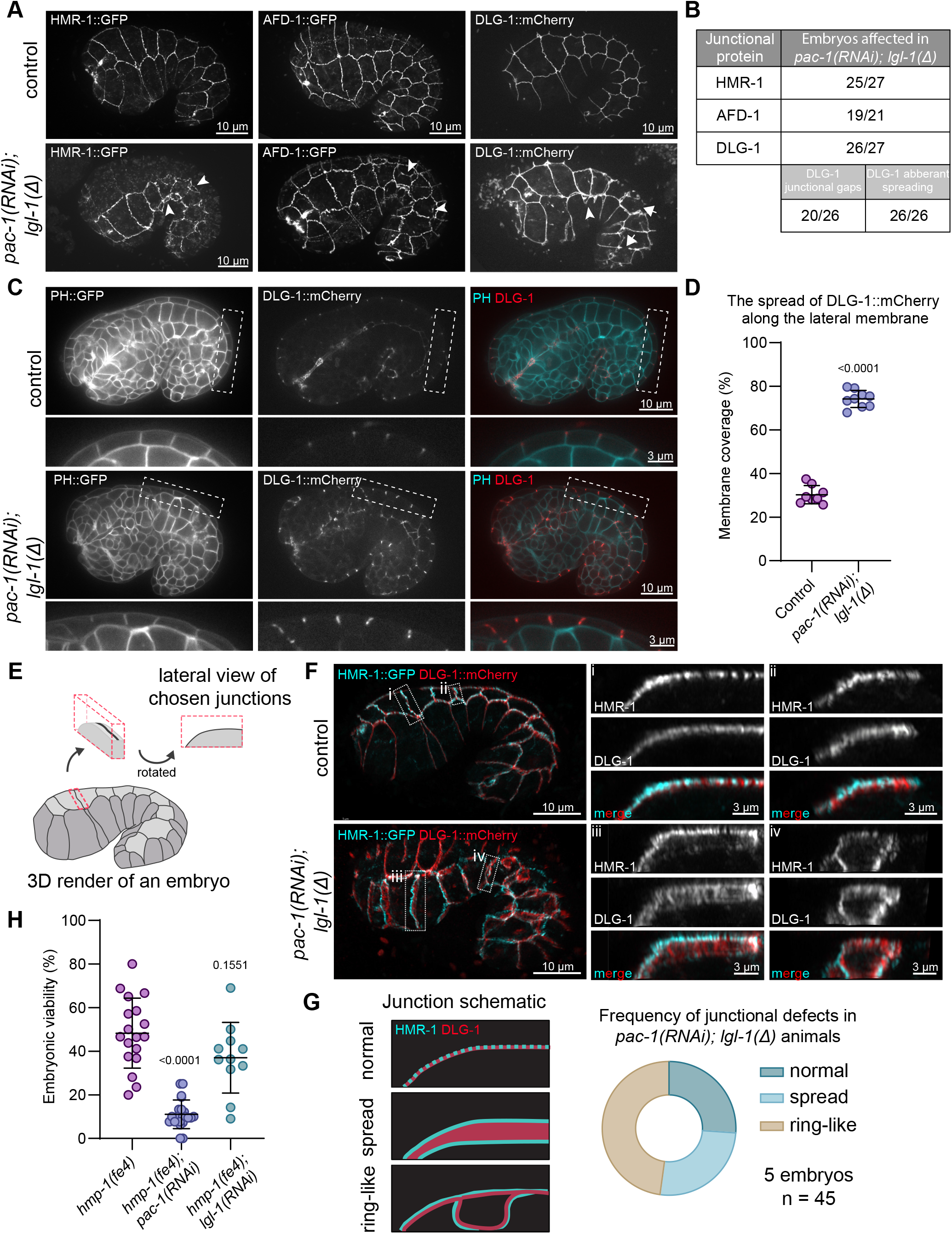
Aberrant localization of junctional marker proteins in *lgl-1(mib201); pac-1(RNAi)* embryos compared to controls. (**A**) Maximum intensity projections of 3D stacks taken with a spinning disc confocal microscope showing the localization of DLG-1::mCherry, AFD-1::GFP and HMR-1::GFP. Arrowheads indicate sites of additional basal junctional signal. Asterisk indicates gap in junctional band. Strains used: BOX804, BOX940, BOX258 and BOX943. (**B**) Quantification of junctional defects. (**C**) Single plane internal view showing the localization of DLG-1::mCherry in respect to the PH::GFP membrane marker. Enlarged versions of the boxed regions are shown below each panel. Images were taken on a spinning disc confocal microscope. Strains used: BOX1076 and BOX1077. (**D**) Quantification of the spread of DLG-1 over the lateral membrane in *pac-1(RNAi); lgl-1(mib201)* mutant animals. Data is represented as mean ± SD and analyzed with Mann-Whitney test; p value shown on graph. Total number of embryos quantified in order of samples in graph: 8 and 9. (**E**) Schematic of the visualization of junctions in panel F. (**F**) 3D rendering of Airyscan confocal images showing DLG-1::mCherry and HMR-1::GFP. Lateral views of the boxed junctions are depicted to the right of the whole embryo images. Strains used: BOX804 and BOX940. (**G**) Schematic of different junctional defects and quantification of their occurrence in *pac-1(RNAi); lgl-1(mib201)* embryos. n = number of junctions quantified. (**H**) Percentage of lethal embryos from *hmp-1(fe4)* and upon knockdown of *lgl-1* or *pac-1*. Data is represented as mean ± SD and analyzed with a Brown-Forsythe and Welch ANOVA test followed by Dunnett’s T3 multiple comparison test; p values shown on graph. Total plates counted in order of samples in graph: 18, 18 and 13. Total progeny counted in order of samples in graph: 514, 500 and 417.

To determine if the defects in junction formation are due to a loss of stability of DLG-1 protein at the membrane, we performed fluorescence recovery after photobleaching (FRAP) experiments using a DLG-1::mCherry endogenous fusion protein. The recovery of DLG-1 was not affected by loss of *lgl-1* and *pac-1*, indicating that the junctional defects are not due to reduced stability of DLG-1 at the membrane (Fig. S2C, D). Furthermore, time-lapse imaging of DLG-1 showed that regions lacking DLG-1 protein remained stable over time (Video S3). Thus, the DLG-1 positioning defects observed in *pac-1; lgl-1* embryos do not appear to stem from a general change in stability of DLG-1 at the membrane, but rather from defects in specifying the positioning of junctional components.

Finally, loss of *pac-1* was previously shown to increase embryonic lethality in the genetically sensitized *hmp-1(fe4)* background. The *fe4* allele encodes a HMP-1^α-catenin^ variant with reduced actin binding that has been used to identify multiple genes that play a role in adherens junction (AJ) functioning (49,51–53). To determine if *lgl-1* may directly affect AJs, we inactivated *lgl-1* in the *hmp-1(fe4)* background. In contrast to *pac-1*, we did not observe any changes in survival of *hmp-1(fe4)* progeny after injection of *lgl-1* dsRNA (Fig. 2H). To rule out that the lack of enhancement by *lgl-1(RNAi)* is due to incomplete inactivation of *lgl-1*, we also re-created the *hmp-1(fe4)* mutation (S823F) by CRISPR in *lgl-1(mib201)* mutant animals and wild-type controls. The phenotype of the S823F mutant we created is more severe than that of the PE97 *hmp-1(fe4)* strain, with only ∼5% of animals becoming fertile adults compared to ∼30 % in PE97 (Fig. S2F) (52). This likely represents the presence of compensatory changes that have accumulated over time in PE97. Nevertheless, consistent with our RNAi results, the presence of *lgl-1(mib201)* did not further exacerbate the phenotype of HMP-1(S823F) (Fig. S2E, F). Taken together, the lack of enhancement of *hmp-1(S823F)* mutants by inactivation of loss of *lgl-1* argues against a primary role for *lgl-1* in regulating cell junctions.

In summary, the combined loss of *lgl-1* and *pac-1* leads to severe defects in the integrity of apical cell junctions, which are likely the cause of the epidermal rupturing and elongation arrest. Unlike *pac-1, lgl-1* does not synergize with junction compromised *hmp-1(fe4)* mutants, indicating that *lgl-1* does not directly regulate AJ formation or functioning.

### Loss of LGL-1 and PAC-1 disrupts apical–basal polarity in the embryonic epidermis

Since both LGL-1 and PAC-1 are thought to directly or indirectly regulate aPKC localization or activity, the junctional defects in *pac-1; lgl-1* embryos may be a result of defects in apical–basal polarization. To investigate this, we examined the localization pattern of aPKC using an endogenous N-terminal GFP::aPKC fusion protein (54). Consistent with previous reports, GFP::aPKC was enriched at the apical cortex and at cell junctions of epidermal epithelial cells during elongation (Fig. 3A). In *pac-1(RNAi); lgl-1(mib201)* embryos, aPKC occupied a broader region apical to cell junctions, indicating enlargement of the apical domain (arrows in Fig. 3A, B and Fig. S3A). Mislocalization of aPKC was especially apparent in lateral regions where DLG-1 was present in a ring-like structure, where aPKC localized in a ring-like structure as well, directly adjacent to or interspersed with the DLG-1 signal (arrowheads in Fig. 3A and Fig. S3A). To determine if increased expression of aPKC might explain the broadened apical localization, we measured total intensity levels of aPKC::GFP. However, we detected no differences in fluorescence levels between control and *pac-1(RNAi); lgl-1(mib201)* animals (Fig. S3B, C).

**Figure 3.**
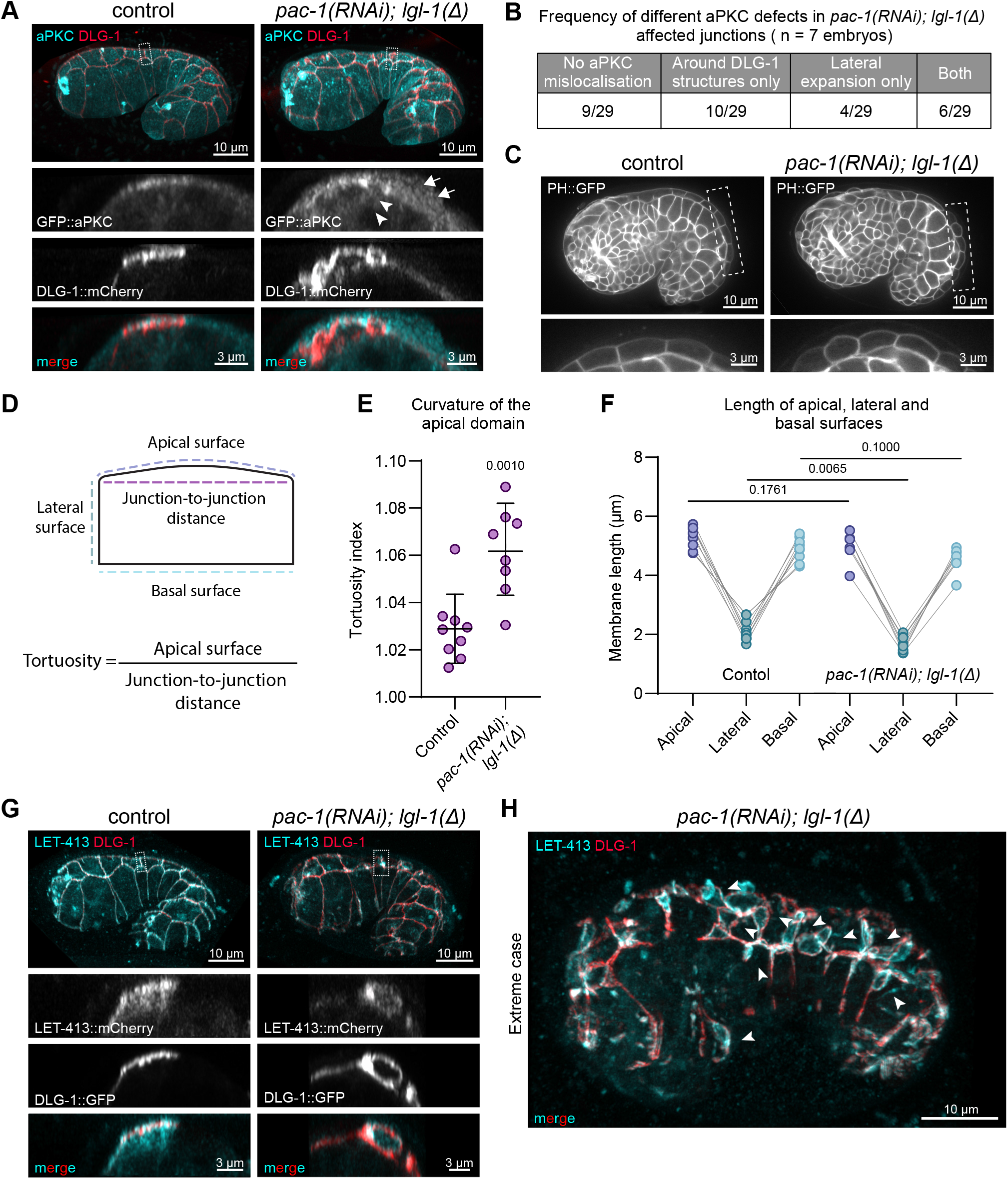
Loss of *lgl-1* and *pac-1* disrupts apical-basal polarity in the embryonic epidermis. (**A**) 3D rendering of Airyscan confocal images showing DLG-1::mCherry and aPKC::GFP. Lateral views of the boxed regions are depicted below the whole embryo images. Asterisk indicates expanded aPKC domain and arrowheads indicate lateral mislocalization of aPKC. Strains used: BOX1161 and BOX1162. (**B**) Qualitative measurement of aPKC mislocalisation in junctions with mislocalised DLG-1::mCherry. n = number of quantified embryos. (**C**) Single plane internal view showing the curvature of the apical domain of the dorsal epidermal cells using the PH::GFP as membrane marker. Enlarged versions of the boxed regions are shown below each panel. Images were taken on a spinning disc confocal microscope. Strains used: BOX1076 and BOX1077. (**D**) Schematic of apical surface tortuosity measurements in Panel E and surface length measurements in Panel F. (**E**) Quantification of the tortuosity of the dorsal epidermal cell apical surface. Each point is the average tortuosity of ≥8 cells in the embryo. Data is represented as mean ± SD and analyzed using the unpaired t test; p value shown on graph. Total number of embryos quantified in order of samples in graph: 9 and 8. (**F**) Quantification of the length of the apical, lateral and basal surface lengths in dorsal epidermal cells. Each point is the average length for ≥8 cells in the embryo. Data is analyzed using the unpaired t test with Welch’s correction; p value shown on graph. Total number of embryos quantified in order of samples in graph: 9 and 8. (**G**) 3D rendering of Airyscan confocal images showing DLG-1::GFP and LET-413::mCherry. Lateral views of the boxed junctions are depicted below the whole embryo images. 7/8 embryos imaged showed one or more junctions with areas devoid of LET-413. Strains used: BOX527 and BOX1100. (**H**) 3D rendering of Airyscan confocal image showing an extreme case of DLG-1::GFP and LET-413::mCherry mislocalization. Arrow heads indicate ring-like distribution of DLG-1 with LET-413 enriched within the rings. Strain used: BOX1100.

Consistent with an expansion of the apical domain, cells appeared more dome shaped than in wild-type controls (Fig. 3B). To quantify this, we determined the tortuosity of the apical surface, defined as the ratio of the length of the cell surface to the direct distance between its end points (see graphic in Fig. 3D). We indeed observed a significantly increased tortuosity of 1.06 compared to 1.03 in controls (Fig. 3E). To better understand the reason for the change in geometry, we also measured the lengths of the lateral and basal surfaces (Fig. 3F). We found that the absolute lengths of the apical surfaces were not significantly different between *pac-1(RNAi); lgl-1(mib201)* and control animals. Instead, the lengths of the lateral domain were reduced (Fig. 3F). Hence, the dome-shaped appearance of epidermal cells in *pac-1; lgl-1* double mutant animals is due to a decrease in lateral domain size, which is consistent with the observed lateral spreading of aPKC.

To further investigate the polarization of epidermal cells in *pac-1; lgl-1* double mutant embryos, we examined the localization of LET-413^Scribble^, which is normally uniformly distributed along the basolateral membrane and enriched at junctions. In *pac-1(RNAi); lgl-1(mib201)* embryos, we observed lateral regions in which LET-413 was absent (Fig. 3G). Strikingly, in all lateral areas with a ring-like distribution of DLG-1, LET-413 was enriched within the ring and absent outside of it (Fig. 3G, H and S3D). Thus, loss of *pac-1* and *lgl-1* can lead to patches of basolateral identity surrounded by junctional material and aPKC.

As aPKC can be recruited by CDC-42, and PAC-1 is a RhoGAP for CDC-42, we also attempted to determine if loss of LGL-1 and PAC-1 affects the activity of CDC-42 using a biosensor consisting of the WSP-1 G-protein binding domain fused to GFP expressed from the cdc-42 promoter (GFP::GBD_WSP-1_) (41,55,56). However, the GFP::GBD_WSP-1_ biosensor was strongly expressed and uniformly localized to the entire cell cortex of embryonic epidermal cells, and therefore not likely to be able to detect the local changes in CDC-42 activity expected from a loss of PAC-1. Indeed, we observed no significant changes in apical GFP::GBD_WSP-1_ membrane levels upon inactivation of lgl-1 and pac-1 (Fig. S3E, F).

Taken together, these data indicate that the combined loss of LGL-1 and PAC-1 results in a shift in the balance of apical vs. basolateral specifying activities towards promotion of apical identity. In most cells, this manifests as an expansion of apical domain identity into the lateral sides of the cells, but in severe cases this can lead to a largely apicalized cell membrane with small patches of basolateral domain (Fig 3E).

### PAC-1 and LGL-1 act in parallel to antagonize aPKC and CDC-42

If the loss of *lgl-1* and *pac-1* leads to a higher level of apical versus basolateral domain specifying activities, then reducing the activity of other apical polarity regulators might partially restore the defects observed in *pac-1; lgl-1* embryos. Two obvious candidates for inactivation are CDC-42 as the presumed target of PAC-1, and aPKC as a major apical domain specifying factor that is counteracted by Lgl.

The complete loss of *cdc-42* leads to embryonic lethality. We therefore partially inactivated *cdc-42* using RNAi by bacterial feeding, which causes only ∼15% lethality by itself (mostly during embryonic development) (Fig. 4A). To avoid complications from performing double RNAi, we examined the effect of *cdc-42* RNAi on the offspring of *pac-1(mib269); lgl-1(mib201)/lgl-1::GFP* parents. Overall, *cdc-42* RNAi caused a mild increase in embryonic viability (Fig. 4A). However, total embryonic viability may underestimate rescue of *pac-1; lgl-1* embryonic lethality, because it also includes the ∼15% lethality caused by *cdc-42* inactivation itself, even among animals wild type for *lgl-1*. We therefore specifically counted animals expressing BFP in the pharynx, which identifies progeny that are homozygous or heterozygous for the *lgl-1(mib201)* deletion. Consistent with our observation that *lgl-1* loss is partially haplo-insufficient, in control populations 45% of viable offspring expressed BFP (as opposed to the expected 67% if *lgl-1* was not haplo-insufficient) (Fig. 4B). On plates with *cdc-42* RNAi bacteria, the fraction of BFP positive animals increased from 45% to 70% (Fig. 4B). Importantly, when examining adult BFP expressing progeny in detail by confocal microscopy, 38/835 did not express any LGL-1::GFP, and were therefore homozygous *pac-1; lgl-1* mutants (Fig. 4C, D). As loss of *pac-1* and *lgl-1* is normally 100% embryonic lethal, this shows that reducing the levels of CDC-42 can partially rescue the embryonic lethality.

**Figure 4.**
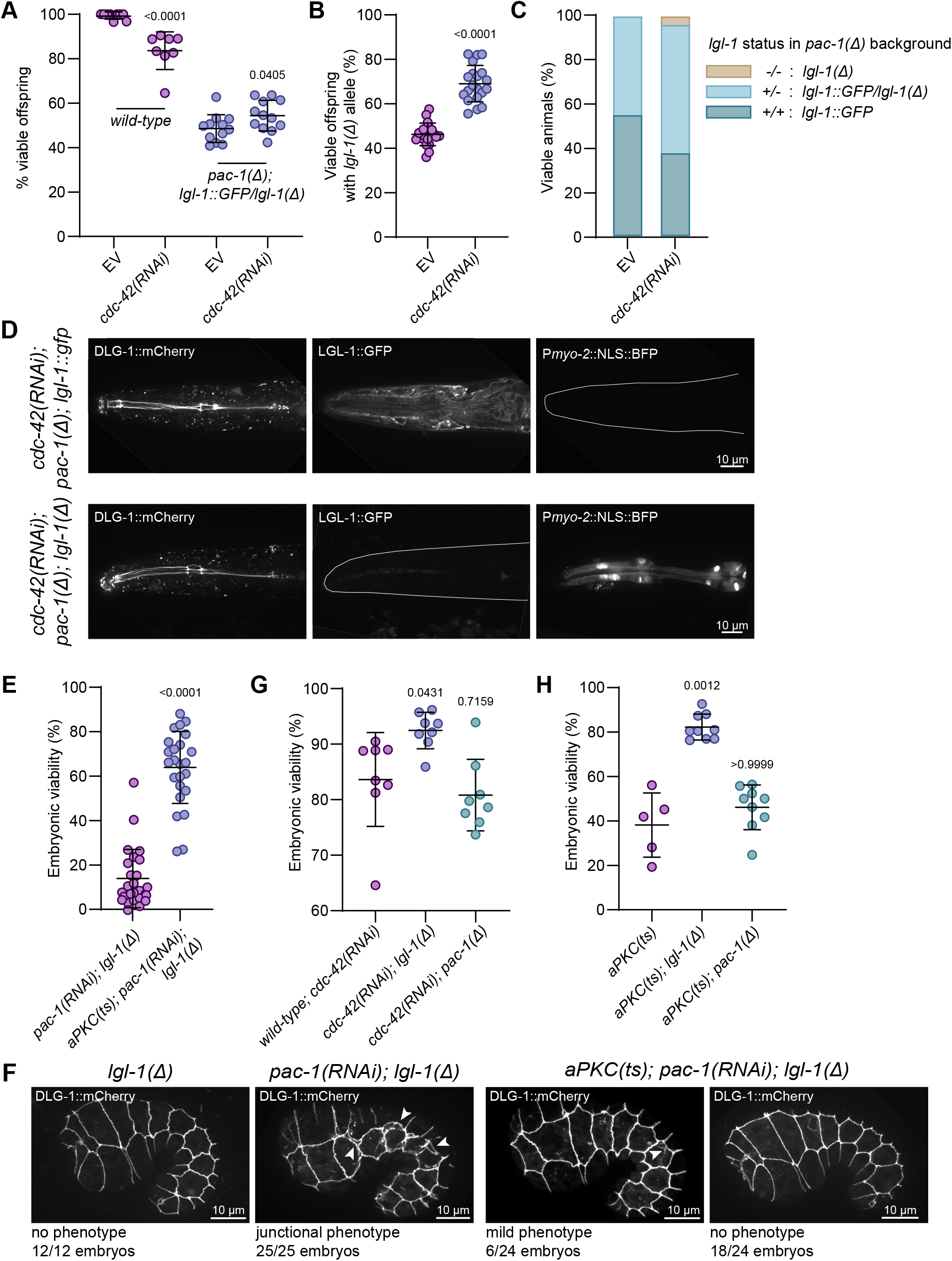
Reduction of CDC-42 or aPKC function rescues the *pac-1; lgl-1* embryonic lethality. (**A**) Percentage of viable wild type and *pac-1(mib269);lgl-1(mib201)/lgl-1::gfp* embryos on empty vector (EV) and *cdc-42* RNAi feeding plates. Data is represented as mean ± SD and the wild type data was analyzed with Mann-Whitney test and the *pac-1(mib269);lgl-1(mib201)/lgl-1::gfp* data was analyzed with unpaired t test with Welch’s correction; p values shown on graph. Total plates counted in order of samples in graph: 9, 8, 12 and 12. Total progeny counted in order of samples in graph: 526, 1068, 1660 and 1627. (**B**) Percentage of the offspring of *pac-1(mib269); lgl-1(mib201)/lgl-1::gfp* animals on either empty vector or cdc-42 RNAi feeding plates, which contain the *lgl-1(mib201)* allele. Data is represented as mean ± SD and analyzed with unpaired t-test with Welch’s correction; p value shown on graph. Total plates counted in order of samples in graph: 20 and 20. Total progeny counted in order of samples in graph: 1132 and 1097. (**C**) Genotype of offspring of *pac-1(mib269); lgl-1(mib201)/lgl-1::gfp* parents on empty vector or *cdc-42* RNAi plates. Total plates counted in order of samples in the graph: 18 and 20. Total progeny counted in order of samples in graph: 906 and 835. (**D**) Single plane images of pharyngeal markers showing examples of two genotypes of animals from the *cdc-42* RNAi plates. Strain used: BOX1176. (**E**) Percentage of viable *pac-1(RNAi); lgl-1(mib201)* embryos in control and upon the partial reduction of aPKC(ts) activity at the semi-permissive temperature of 20°C. Data is represented as mean ± SD and analyzed with Mann-Whitney test; p value shown on graph. Total plates counted in order of samples in the graph: 26 and 26. Total progeny counted in order of samples in graph: 2244 and 1995. (**F**) Maximum intensity projections of 3D stacks taken with a spinning disc confocal microscope showing the localization of DLG-1::mCherry. Arrowheads indicate ectopic basal junctional signals. Strains used: BOX1179 and BOX1177. (**G**) Percentage of viable embryos from wild-type, *lgl-1(mib201)* or *pac-1(mib269)* on *cdc-42* RNAi. Data is represented as mean ± SD and analyzed with Kruskal-Wallis statistical test with Dunn’s multiple comparison test; p values shown on graph. Total plates counted in order of samples in the graph: 8, 8 and 8. Total progeny counted in order of samples in graph: 1068, 916 and 958. (**H**) Percentage of viable embryos from *aPKC(ts), aPKC(ts); lgl-1(mib201)* or *aPKC(ts); pac-1(mib269)*, at semi-permissive temperature of 20°C. Data is represented as mean ± SD and analyzed with Kruskal-Wallis statistical test with Dunn’s multiple comparison test; p values shown on graph. Total plates counted in order of samples in the graph: 5, 9 and 9. Total progeny counted in order of samples in graph: 386, 792 and 612.

Like CDC-42, the complete loss of aPKC is embryonic lethal. To investigate the effect of reducing aPKC activity on animals lacking LGL-1 and PAC-1, we therefore used the temperature sensitive allele a*PKC(ne4250)*, which ontains an H390Q in the kinase domain (57). We generated an a*PKC(ts); lgl-1(mib201)* double mutant strain and inactivated *pac-1* by dsRNA injection. The reduced aPKC activity at the semi-permissive temperature of 20°C significantly restored embryonic viability of *pac-1(RNAi); lgl-1(mib201)* animals, from 17% in controls to 63% (Fig. 4E). We also investigated the appearance of cell junctions in a*PKC(ts); pac-1(RNAi); lgl-1(mib201)* animals in a strain expressing DLG-1::mCherry as well as an endogenous PAC-1::GFP fusion protein that labels all PAC-1 isoforms (Fig. S4A). At the semi-permissive temperature, the reduced aPKC activity rescued the junctional defects of the *pac-1; lgl-1* mutant animals (Fig. 4F). Moreover, the presence of PAC-1::GFP allowed us to rule out that the rescue by *aPKC(ts)* is due to an effect on *pac-1* RNAi efficiency, as we observed the expected strong reduction of PAC-1::GFP signal (Fig. S4B).

The rescue of the combined loss of PAC-1 and LGL-1 by reducing the function of CDC-42 or aPKC is consistent with PAC-1 and LGL-1 acting in parallel pathways that both regulate aPKC activity. We therefore wanted to investigate the relative contributions of PAC-1 and LGL-1 to the regulation of aPKC. To accomplish this, we determined whether the loss of PAC-1 or LGL-1 individually could compensate for the embryonic lethality associated with partial loss of aPKC or CDC-42. Strikingly, loss of *lgl-1* increased the viability of both *cdc-42(RNAi)* and *aPKC(ts)* embryos, but the loss of *pac-1* had no significant effect (Fig. 4G, H).

Taken together our findings support a model in which PAC-1 and LGL-1 act in parallel to antagonize the apical polarizing activities of aPKC and CDC-42. Since loss of individual LGL-1, but not of PAC-1, also suppresses the embryonic lethality of *cdc-42(RNAi)* or *aPKC(ts)* embryos, LGL-1 appears to play a stronger role in the network of inhibitory interactions that establish apical–basal polarity than PAC-1, while PAC-1 may play a more local role at cell junctions.

## Discussion

Here we show that LGL-1 acts redundantly with the RhoGAP protein PAC-1 to control morphogenesis of the embryonic epidermal epithelium of *C. elegans*. Loss of *lgl-1* alone does not cause defects in cell polarization and does not affect viability. Loss of *pac-1* causes defects in cell positioning during gastrulation, but ∼90% of embryos recover and hatch with no morphological defects (39,41). In contrast, the combined loss of *pac-1* and *lgl-1* leads to a fully penetrant lethal phenotype. Most animals arrest as embryos around the 2-fold stage of elongation with severe morphological defects in the epidermis, including rupture. These defects are likely caused by a failure to properly polarize epidermal epithelial cells, as we observed an increased apical domain size, defects in the localization of basolateral LET-413^Scribble^, and severe defects in the positioning of cell junctions.

The severe defects in apical–basal polarization strongly suggest that PAC-1 and LGL-1 act redundantly to control epithelial polarity. At what level do these two proteins act redundantly? A likely candidate is the regulation of aPKC. In flies and mammalian model systems, antagonism between Lgl and aPKC is well established (1–4). Previous studies in *C. elegans* have implicated LGL-1 as a regulator of aPKC in the zygote and spermatheca. Loss of LGL-1 in one-cell embryos that also lack PAR-2 results in an expansion of the anterior PAR-6–aPKC domain (35,36), and *lgl-1* loss can suppress the sterility observed in a temperature sensitive *aPKC* mutant, which is likely due to junctional defects in the spermatheca (38). Our genetic analysis now indicates that LGL-1 acts as an inhibitor of aPKC activity in the epidermal epithelium. First, the combined loss of PAC-1 and LGL-1 leads to increased apical domain size at the expense of the basolateral domain, phenotypes consistent with overactivation of aPKC. Second, the lethality of combined loss of *pac-1* and *lgl-1* can be suppressed by reducing aPKC activity. Finally, loss of *lgl-1* alone suppresses the embryonic lethality of *aPKC(ts)* mutants at the semi-permissive temperature. While additional non-aPKC dependent roles are possible, the predominant function of LGL-1 in the epidermis therefore likely is the inhibition of aPKC. Thus, *C. elegans* LGL-1 has now been shown to act as an inhibitor of aPKC in multiple tissues.

PAC-1 was also identified as a regulator of aPKC during radial polarization of *C. elegans* blastomeres. In *pac-1* mutant blastomere stage embryos, PAR-6 and aPKC localize uniformly to the entire cell cortex rather than being enriched at the contact-free apical surface (39). Experiments using constitutively active CDC-42 and PAR-6 mutants unable to bind to CDC-42 indicate that PAC-1 locally inactivates CDC-42 at cell–cell contacts to restrict PAR-6–aPKC recruitment to the apical contact-free surface (39). Inhibition of aPKC activity via downregulation of Cdc42 is further supported by studies of the *Drosophila* PAC-1 ortholog RhoGAP19D. Similar to *C. elegans* blastomeres, RhoGAP19D is recruited to E-cadherin adhesion complexes in the *Drosophila* follicular epithelium, and *rhogap19d* mutants show increased lateral Cdc42 activity together with an expansion of the apical domain due to higher Par6–aPKC activity (42). Moreover, *rhogap19d* mutant follicle cells invade the overlying germline cyst. This phenotype is suppressed by loss of one copy of *aPKC*, and enhanced by loss of one copy of *lgl*, consistent with our genetic analyses (42).

Interestingly, the roles of PAC-1 and CDC-42 have previously been investigated in the embryonic epidermis. In that study, PAC-1 and CDC-42 were shown to be involved in regulating junctional actin dynamics and adherens junction protein levels during elongation, but to be dispensable for polarization and junction maturation (41). Our data examining the roles of PAC-1 and CDC-42 in an *lgl-1* mutant background now demonstrate that both proteins are important regulators of epidermal cell polarity, consistent with the previously described function of PAC-1 in *C. elegans* blastomeres (39), the essential role of CDC-42 regulating aPKC activity in the *C. elegans* zygote (11), and the roles of orthologs of both proteins in other systems. Whether the previously observed defects in actin regulation and adherens junction protein levels reflect aPKC independent functions of CDC-42 and/or PAC-1 or are a manifestation of underlying minor polarity defects remains to be determined. While the defects in polarization and junction establishment are severe, we cannot exclude therefore that non-aPKC dependent roles of CDC-42 and/or PAC-1 contribute to the morphogenesis defects of *pac-1 lgl-1* double mutants.

Taken together, the simplest model integrating our findings is that analogous to recent models proposed for *Drosophila*, PAC-1 and LGL-1 act in parallel pathways that oppose aPKC activity (Fig 5). In this model, LGL-1 directly binds to and inhibits PAR-6/aPKC, while PAC-1 indirectly reduces aPKC activity by locally inactivating CDC-42. While the overall mechanisms of epithelial polarization are thus conserved between *C. elegans* and *Drosophila*, important differences still exist. In *C. elegans*, neither LGL-1 nor PAC-1 are essential proteins, whereas Lgl and RhoGAP19D are both individually required for epithelial polarization (24,42,58). Another difference is the importance of the Crumbs proteins. While the single *Drosophila* Crumbs protein is essential for epithelial polarization (59,60), a triple deletion strain lacking all three *C. elegans crumbs* genes is viable and shows no polarity defects (33). Nevertheless, minor defects in epithelial organization upon loss of the Crumbs protein CRB-1 can been observed in a mutant background lacking the Scribble ortholog *let-413* and the α-catenin ortholog *hmp-1* (61), and overexpression of the Crumbs protein EAT-20 or CRB-3 in the intestine causes mild enlargement of the apical domain (32). One interpretation of these differences is that, for reasons remaining to be discovered, regulation of aPKC is more critical in *Drosophila* than in *C. elegans*. Alternatively, the strength of aPKC regulation by RhoGAP19D, Lgl, and Crumbs may be stronger in *Drosophila* than in *C. elegans*, such that loss of any regulator more strongly affects overall aPKC activity levels.

**Figure 5.**
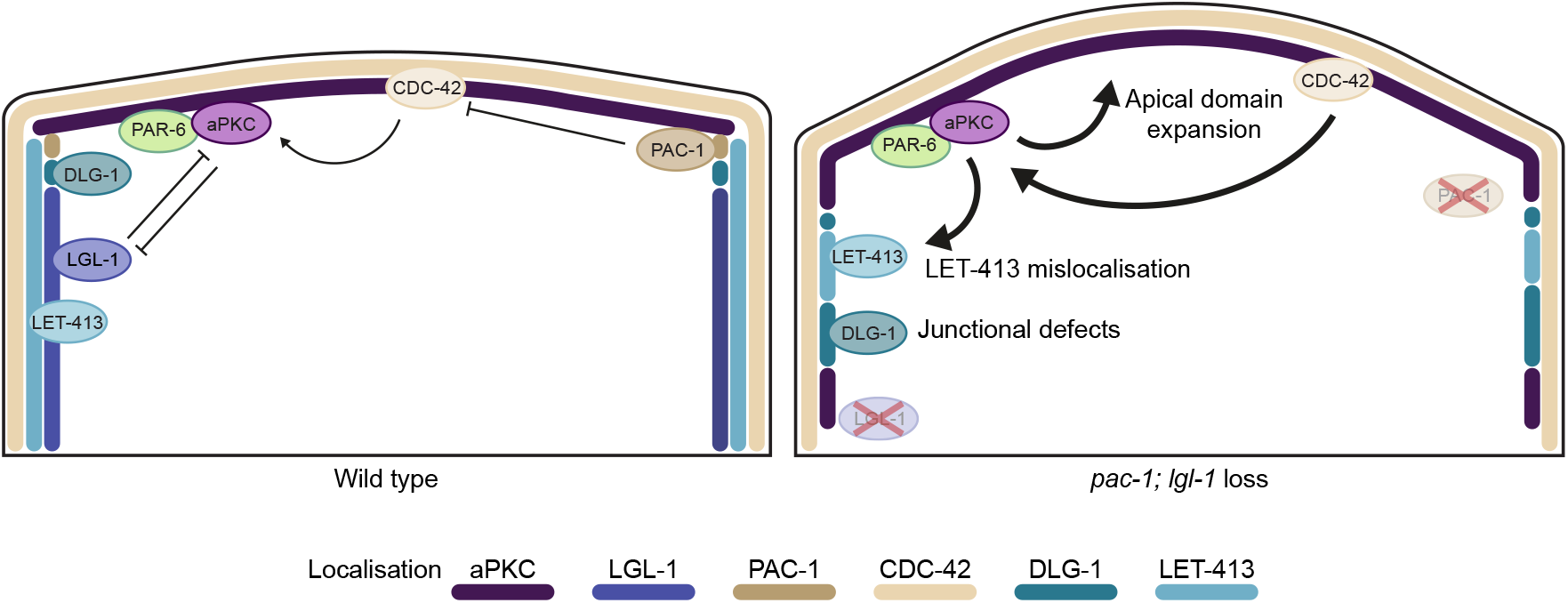
Model of LGL-1 and PAC-1 function in the *C. elegans* embryonic epidermal cells. Colored lines indicate the relative localization of proteins.

Redundancy between Lgl and other polarity proteins may not be unique to *C. elegans*. The role of Lgl proteins as negative regulators of aPKC is well established in mammalian models (8,9,62–65). However, whether Lgl is broadly essential for polarity establishment in mammals has been difficult to assess due to the presence of two paralogs, individual knockouts of which cause tissue-specific and late appearing phenotypes. In mice, knockout of Lgl1 specifically affects polarization of neuronal cells, and knockout mice die shortly after birth from severe hydrocephalus (64), while Lgl2 knockout mice are born small due to a placental branching defect but develop into normal adults (66). In zebrafish, only cell-type specific roles have been reported upon Lgl loss, including neuroepithelial polarity defects for Lgl1 (67) and basal and peridermal formation defects for Lgl2 (65,68,69). The presence of two paralogs may be an explanation for the lack of more severe defects. However, a recent preprint study generating conditional double knockouts found that mice lacking both *Lgl1* and *Lgl2* in the skin epidermis are viable and fertile (70). Thus, while the complete absence of polarity phenotypes upon Lgl loss appears unique to *C. elegans*, Lgl proteins may well act redundantly with other polarity pathways in vertebrates as well.

Finally, while we favor a model based on regulation of aPKC activity by PAC-1 via CDC-42, other interpretations are possible. In the epidermis, degradation of CDC-42 does not block polarization (41). While apical PAR-6 levels are reduced, PAR-6 is still asymmetrically localized in CDC-42 depleted embryos. Thus, additional mechanisms to regulate PAR-6/aPKC localization may remain to be discovered. A recent study of blastomere stage polarization has also suggested that PAC-1 may regulate aPKC independently of CDC-42 (71). Quantitative analysis of aPKC levels showed that the uniform localization of aPKC in *pac-1* mutants is due to a loss of apical aPKC rather than to ectopic lateral recruitment. More importantly, CDC-42 degradation using the ZF1 degron did not prevent aPKC polarization, yet in this CDC-42 depleted background *pac-1* RNAi still caused aPKC to become uniformly localized. These results are inconsistent with PAC-1 acting via CDC-42. As PAC-1 can act as a GAP for other Rho-family small GTPases (72), an alternative hypothesis that was suggested is that PAC-1 acts via a different small GTPase. Our own genetic data cannot elucidate whether PAC-1 acts solely via CDC-42 and aPKC. While the rescue of embryonic lethality of *pac-1 lgl-1* double mutants by partial loss of *cdc-42* or *aPKC* is consistent with this, it was surprising to us that loss of *pac-1* alone did not affect embryonic lethality of *cdc-42(RNAi)* or *aPKC(ts)* animals. This result could be interpreted as PAC-1 not acting in the same pathway as either CDC-42 or aPKC. However, even if PAC-1 were to act in a separate pathway controlling apical–basal polarity, it’s loss would still be expected to affect the phenotypes of other mutants in polarity regulators, including CDC-42 or aPKC.

One difficulty in resolving the precise relationship between PAC-1, CDC-42, and aPKC is that we cannot directly assess the activity of aPKC, and a currently available CDC-42 biosensor is not very sensitive to small or local changes in CDC-42 activity, due to its strong and uniform cortical localization. Only severe reductions in CDC-42 activity, *e*.*g*. through *cdc-42(RNAi)* or overexpression of PAC-1, have been reported to result in consistent changes in localization of the biosensor (41,56). In *Drosophila*, analysis of Cdc42 sensors more clearly support that RhoGAP19D acts via Cdc42, as RhoGAP19D loss results in lateral recruitment of the Cdc42 effectors N-WASP and Gek (42). Moreover, induced recruitment of RhoGAP19D to the apical domain mimics *cdc42* mutants. For these reasons, we consider it likely that PAC-1 does act (predominantly) via CDC-42 and aPKC but plays a smaller role in regulating aPKC activity than LGL-1. As long as LGL-1 is present, the loss of PAC-1 has little effect on aPKC activity.

In summary, our study shows that the overall network of proteins regulating epithelial polarity is highly conserved in *C. elegans*. The surprising finding that LGL-1 is a non-essential protein does not reflect an alternative mode of polarization but likely reflects variations in the strengths of connections within this polarity network.

## Materials and Methods

### *C. elegans* strains and culture conditions

*C. elegans* strains were cultured under standard conditions (73). Only hermaphrodites were used and all experiments, until stated otherwise, were performed with animals grown at 15°C on Nematode Growth Medium (NGM) agar plates seeded with *Escherichia coli* OP50 bacteria. Table 1 contains a list of all the strains used. All strains were derived from N2/CGC1.

**Table 1.** List of *C. elegans* strains used in the study.

| Designation | Genotype | Source or reference |
| --- | --- | --- |
| CGC1 | wild-type | CGC |
| FT1459 | *xnIs506 [Pcdc42::GST::GFP::wsp-1(GBD) + unc-119(+)]* | (41) |
| PE97 | *hmp-1(fe4[S823F]) V* | (53) |
| NWG0285 | *lgl-1(crk66[lgl-1::gfp]) X* | (79) |
| WM151 | *pkc-3(ne4250 [E9K; H390Q]) II* | Gift from Rodriguez lab |
| BOX258 | *afd-1(mib34[afd-1::GFP-LoxP]) I* | This study |
| BOX315 | *lgl-1(mib44[Pmyo-2::GFP, X:863693..874171]) X* | This study |
| BOX443 | *pkc-3(mib78[egfp-loxp::aid::pkc-3])II* | (54) |
| BOX527 | *mibIs49[Pwrt-2::TIR-1::tagBFP2-LoxP::tbb-2-3'UTR, IV:5014740-5014802 (cxTi10816 site))] IV; mib29[let-413::mCherry-LoxP]) V; dlg-1(mib35[dlg-1::AID::GFP-LoxP]) X* | (80) |
| BOX804 | *hmr-1(he298[hmr-1::GFP]) I; mibIs49[Pwrt-2::TIR-1::tagBFP2-LoxP::tbb-2-3'UTR, IV:5014740..5014802 (cxTi10816 site))] IV; dlg-1(mib128[dlg-1::AID::mCherry-LoxP]) X* | This study |
| BOX899 | *lgl-1(mib201[Pmyo-2::NLS::mTagBFP2, X:863693..874171]) dlg-1(mib23[dlg-1::mCherry-LoxP]) X* | This study |
| BOX907 | *pac-1(mib269[III: 8494328..8519244]) III* | This study |
| BOX919 | *lgl-1(mib201[Pmyo-2::NLS::mTagBFP2, X:863693..874171]) X* | This study |
| BOX940 | *hmr-1(he298[hmr-1::GFP]) I; lgl-1(mib201[Pmyo-2::NLS::mTagBFP2, X:863693..874171]) dlg-1(mib128[dlg-1::AID::mCherry-LoxP]) X* | This study |
| BOX943 | *afd-1(mib34[afd-1::GFP-LoxP]) I; lgl-1(mib201[Pmyo-2::NLS::mTagBFP2, X:863693..874171]) X* | This study |
| BOX992 | *xnIs506 [Pcdc42::GST::GFP::wsp-1(GBD) + unc-119(+)]; lgl-1(mib-201[Pmyo-2::NLS::mTagBFP2, X:863693..874171]) X* | This study |
| BOX1027 | *pkc-3(ne4250 [E9K; H390Q]) II; lgl-1(mib201[Pmyo-2::NLS::mTagBFP2, X:863693..874171]) X* | This study |
| BOX1045 | *pkc-3(mib78[egfp-loxp::aid::pkc-3])II; lgl-1(mib201[Pmyo-2::NLS::mTagBFP2, X:863693..874171]) X* | This study |
| BOX1075 | *pac-1(mib296[pac-1::mIAA7::GFP]) III* | This study |
| BOX1076 | *cxTi10816(he259[Peft-3::PH::GFP::LOV::tbb-2]); dlg-1(mib-23[dlg-1::mCherry-LoxP]) X* | This study, PH::GFP from (81) |
| BOX1077 | *cxTi10816(he259[Peft-3::PH::GFP::LOV::tbb-2]); lgl-1(mib-201[Pmyo-2::NLS::mTagBFP2, X:863693..874171]) dlg-1(mib-23[dlg-1::mCherry-LoxP]) X* | This study |
| BOX1100 | *let-413 (mib29[let-413::mCherry-LoxP]) V/+; lgl-1(mib201[Pmyo-2::NLS::mTagBFP2, X:863693..874171]) dlg-1(mib35[dlg-1::AID::GFP-LoxP]) X* | This study |
| BOX1102 | *pac-1(mib296[pac-1::mIAA7::GFP])III;*<br>*lgl-1(mib201[Pmyo-2::NLS::mTagBFP2, X:863693..874171]) X* | This study |
| BOX1106 | *lgl-1(crk66[lgl-1::gfp]) X;*<br>*pac-1(mib269[III: 8494328..8519244]) III* | This study |
| BOX1111 | *pac-1(mib269[III: 8494328..8519244]) III; lgl-1(crk66[lgl-1::GFP])*<br>*/lgl-1(mib201[Pmyo-2::NLS::mTagBFP2, X:863693..874171]) X* | This study |
| BOX1161 | *pkc-3(mib78[GFP-LoxP::AID::pkc-3]) II; lgl-1(mib201[Pmyo-2::N-*<br>*LS::mTagBFP2, X:863693..874171]) dlg-1(mib23[dlg-1::mCher-*<br>*ry-LoxP]) X* | This study |
| BOX1162 | *pkc-3(mib78[GFP-LoxP::AID::pkc-3]) II; dlg-1(mib23[dlg-1::mCher-*<br>*ry-LoxP]) X* | This study, *dlg-1* from (82) |
| BOX1176 | *pac-1(mib269[III: 8494328..8519244]) III; lgl-1(crk66[lgl-1::G-*<br>*FP])/lgl-1(mib201[Pmyo-2::NLS::mTagBFP2, X:863693..874171])*<br>*dlg-1(mib23[dlg-1::mCherry-LoxP])/+ X* | This study |
| BOX1177 | *pkc-3(ne4250 [E9K; H390Q]) II; pac-1(mib296[pac-1::mIAA7::G-*<br>*FP])III; lgl-1(mib201[Pmyo-2::NLS::mTagBFP2,*<br>*X:863693..874171]) X dlg-1(mib23[dlg-1::mCherry-LoxP]) X* | This study |
| BOX1179 | *pac-1(mib296[pac-1::mIAA7::GFP]) III; lgl-1(mib201[Pmyo-2::N-*<br>*LS::mTagBFP2, X:863693..874171]) dlg-1(mib23[dlg-1::mCher-*<br>*ry-LoxP]) X* | This study |
| BOX1195 | *pkc-3(ne4250 [E9K; H390Q]) II; pac-1(mib269[III:*<br>*8494328..8519244]) III* | This study |
| BOX1221 | *pac-1(mib269[III: 8494328..8519244]) III; lgl-1(mib44[Pmyo-2::G-*<br>*FP, X:863693..874171]) X /+* | This study |
| BOX1344 | *hmp-1(mib387[S823F]) V; lgl-1(mib201[Pmyo-2::NLS::mTagBFP2,*<br>*X:863693..874171]) X* | This study |
| BOX1348 | *hmp-1(mib386[S823F]) V* | This study |

### CRISPR

The *lgl-1(mib44)* knockout allele and *afd-1(mib34)* knock-in allele, were generated using plasmid-based expression of Cas9 and sgRNA using the self-excising cassette (SEC) for selection (74) and the *lgl-1(mib201), pac-1(mib269), pac-1(mib296)* and *dlg-1(mib128)* were generated using a plasmid-free approach (75). The *lgl-1(mib44)* edit was made in N2 background and the other edits were made in the CGC1 background. In cases where the sgRNA target site was not disrupted by sequence integration, silent mutations were incorporated to prevent repeated DNA cleavage. The sequences to generate the CRISPR alleles are found in table 2. The full genome sequences of the final strains are available in Supplementary File 1.

**Table 2.**
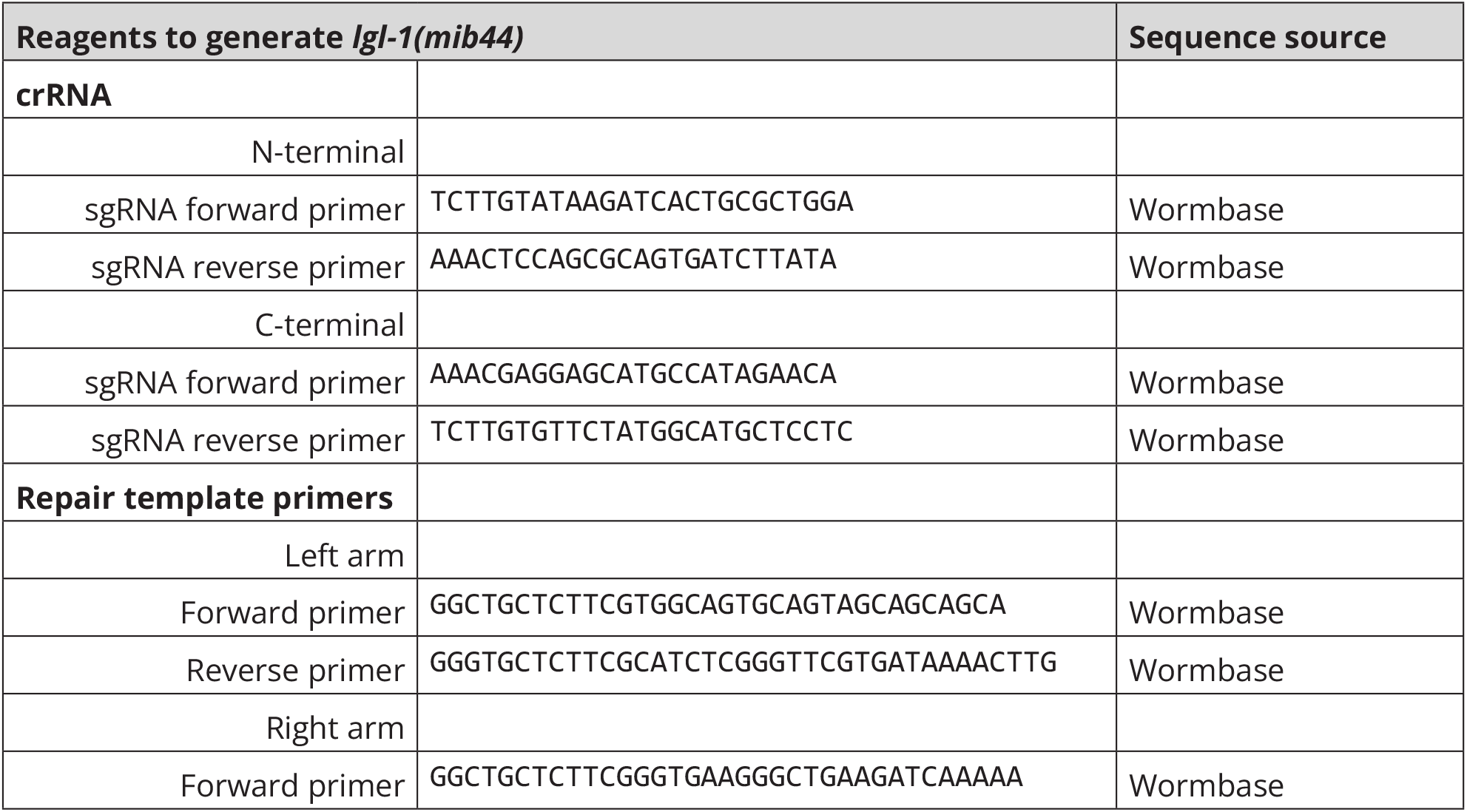
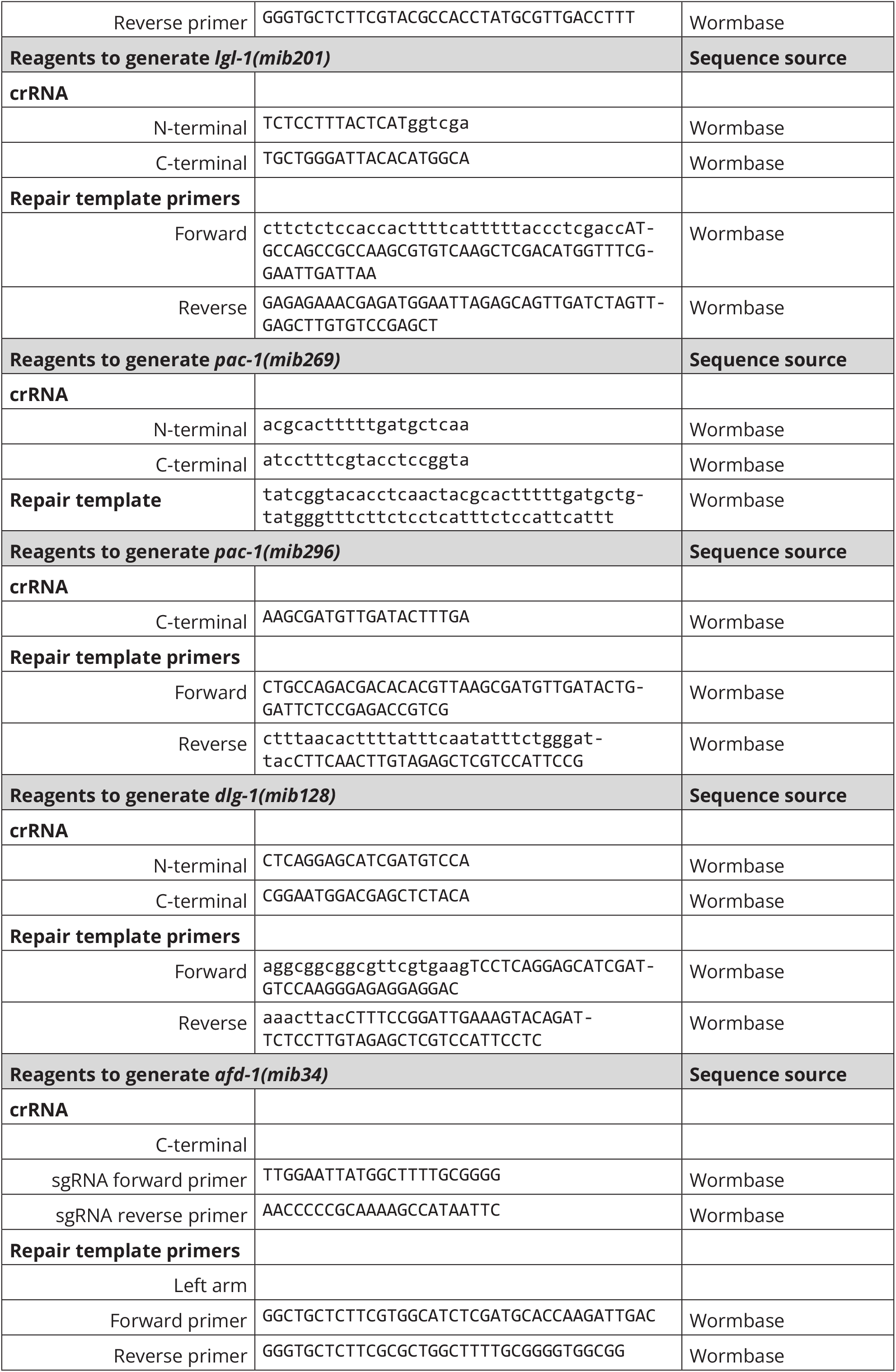
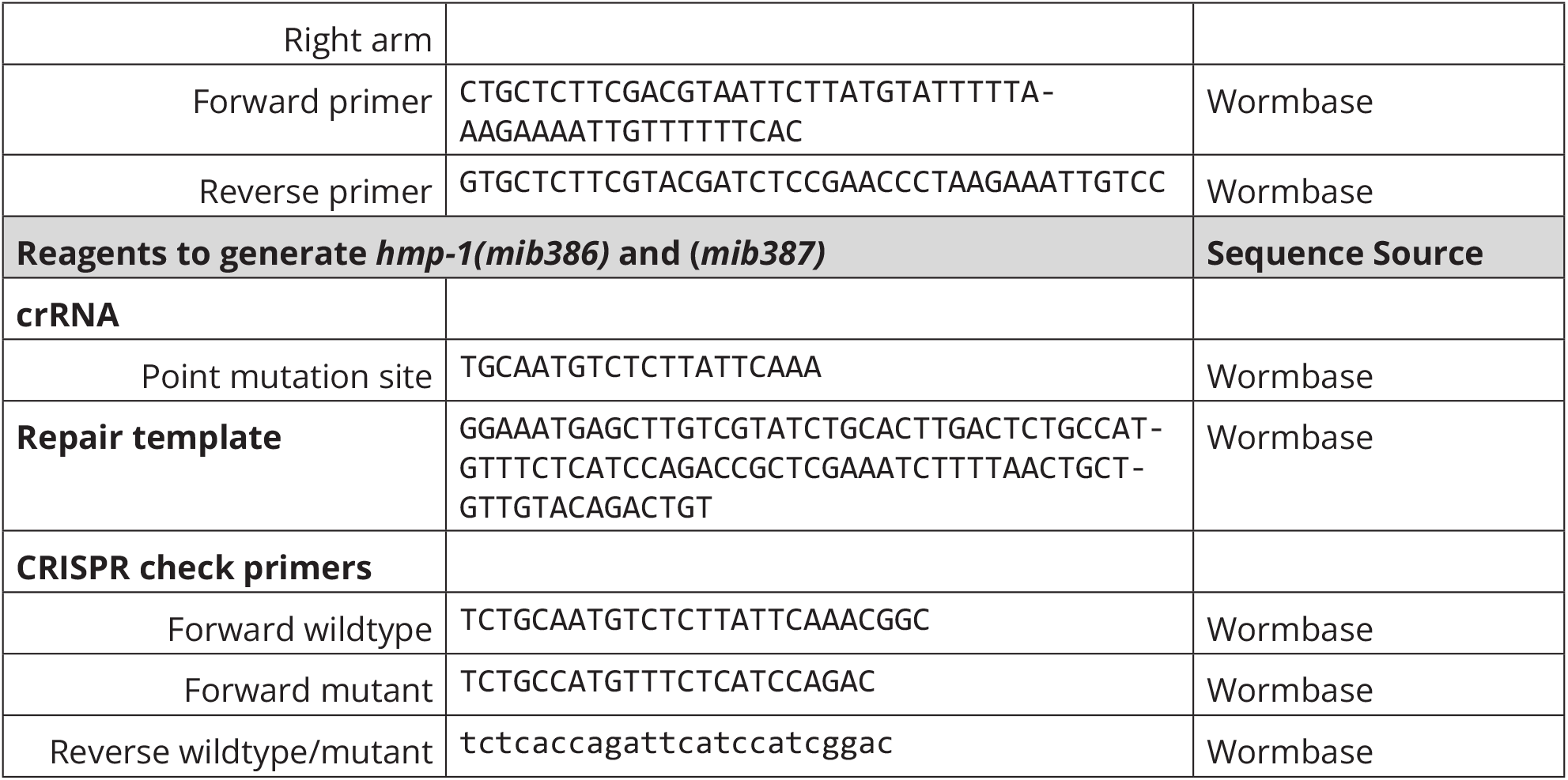
List of guide RNA and primers used to generate the CRISPR edited animals.

For the plasmid-based CRISPR/Cas9 editing, the homology arms for *lgl-1(mib44)* were cloned in SapTrap assembly vector pMLS257 (74). The sgRNAs were expressed from a plasmid under a control of a U6 promoter. To generate sgRNA vectors, antisense oligonucleotide pairs were annealed and ligated into Bbsl-linearized pJJR50 (Addgene ID #75026) (76). Injection mixes were prepared in MilliQ water and contained 50 ng/ml *Peft-3::cas9* (Addgene ID #46168) (77) 50-100 ng/ml *U6::sgRNA*, and 50-75 ng/ml of repair template. The mix contained 2.5 ng/ml of the co-injection pharyngeal marker *Pmyo-2::tdTomato* to aid in the visual selection of transgenic strains. Young adult hermaphrodite germlines were microinjected using an inverted microinjection setup (Eppendorft FemtoJet 4x mounted on a Zeiss Axio Observer A. 1 equipped with an Eppendorf Transferman 4r). The candidate edited progeny were selected on plates contain 250 ng/ml of hygromycin (74) and correct genome editing was confirmed by Sanger sequencing (Macrogen). From the correctly edited animals, the hygromycin selection cassette was excised by a heat shock of L1 larvae at 34°C for 1 hour in a water bath.

For the knock-in and knockout strains generated using the plasmid-free approach, the repair templates were PCR amplified using Q5 Hot Start Polymerase (75). The amplification of the repair template was done from the following pre-existing donor vectors: GFP from pJRK86 (Addgene ID #173743), BFP from pDD356, and mCherry from pRS004. The generated animals were sequenced for correct gene editing by Sanger sequencing (Macrogen).

### Genome wide-RNAi screen

The RNAi screen was conducted as described in (43). The administered RNAi feeding used the publicly available Ahringer RNAi clone library (44). The primary screen was conducted using the NYUAD high-throughput screening automated platform, using the wild-type N2 and *lgl-1(mib44)* animals, in two replicates (described in (78)). From the screen, 79 clones caused lethality, developmental delay or arrest, sterility, embryonic lethality or morphological defects in the *lgl-1(mib44)* animals and not or to a lesser extent in the wild-type. 27 of the clones were discarded as false positives, as they are frequently found in other RNAi screens done using the same pipeline. The rest of the 52 clones were independently tested by putting L1 animals on RNAi feeding using sequence verified Ahringer RNAi library clones (see section RNAi treatment).

### RNAi treatment

All RNAi clones for the verification screen and targeted experiments were obtained from the genome wide Ahringer HT115 RNAi feeding library, supplied through Source BioScience, and verified using Sanger sequencing.

For feeding RNAi experiments, bacteria were pre-cultured in 2–4 ml lysogeny broth (LB) supplemented with 100 µg/ml ampicillin (Amp) and 2.5 µg/ml tetracycline (Tet) at 37°C in an incubator rotating at 220 rpm. Bacteria were then diluted 1:5 in 10 ml LB Amp/Tet for the genome-wide RNAi verification screen or 1:1000 in 25 ml LB Amp/Tet for the *cdc-42* RNAi feeding experiments. To induce production of dsRNA, the cultures were incubated for an hour with 1mM Isoporyl b-D-1-thiogalactopyranoside (IPTG). Bacterial cultures were pelleted by centrifugation at 4000 g for 15 minutes at 4°C, and concentrated 5 times. NGM plates were supplemented with 100 μg/ml Amp, 2.5 µg/ml Tet and 1 mM IPTG, seeded with 200 µl of bacteria, kept at room temperature for 48 hours in the dark, until dry then kept at 4°C until used.

For the *cdc-42* RNAi experiments, young adults were placed on the seeded RNAi plates for 2 days, at 20°C. The following day the number of animals with a visible *lgl-1(mib201)* pharyngeal marker were counted and scored for their genotype based on their fluorescence, using a spinning disc fluorescence microscope.

For stronger RNAi, young adults were injected with dsRNA targeting *pac-1* and *lgl-1*. To generate dsRNA, *pac-1* and *lgl-1* clones from the Ahringer RNAi library were amplified using Q5 Hot Start High-Fidelity DNA polymerase (New England Biolabs). The PCR product was used as a template for in vitro dsRNA synthesis using MEGAscript T7 transcription kit (ThermoFisher Scientific). dsRNA was diluted 1:5 in RNase free water prior to micro-injection, and kept at -80°C. The injected animals were kept at 15°C for 24 hours, then transferred to new plates and used for lethality assays.

### Embryonic viability and lethality assays

Young adult animals were cultured at 15°C on OP50 plates, and allowed to lay eggs for 24–36 hours. Animals were subsequently removed and after 24 hours the number of hatched or unhatched eggs was counted. Embryos that did not hatch and young dead larvae were counted as dead. For *aPKC(ts)* experiments young adults were allowed to lay eggs at the permissive temperature of 15°C for 20 hours, then were moved to a semi-permissive temperature of 20°C for 8 hours. The adults were then taken off and the embryos were left to hatch at 20°C for 24 hours before counting viability.

### Imaging

Imaging of *C. elegans* was done by mounting embryos or larvae on a 5% agarose pad in a 10 mM Tetramizole solution in M9 buffer to induce paralysis. DIC imaging was performed using an upright Zeiss Axio lmager Z2 microscope using a 100x 1.4 NA objective and Zeiss AxioCam 503 monochrome camera, driven by Zeiss Zen Blue Software version 3.3. Spinning disc confocal imaging was performed using a Nikon Ti-U microscope driven by MetaMorph Microscopy Automation and Image Analysis Software version 7.10.5.476 (Molecular Devices) and equipped with Yokogawa CSU-X1-M1 confocal head and a Prime BSI sCMOS camera (Teledyne Photometrics), using Nikon CFI Plan Apo λ 60 x or 100 × 1.4 NA objectives. Imaging on the Zeiss AxioCam 503 and Nikon Ti-U was done in a temperature controlled room, at 20°C. AiryScan imaging was done using Confocal Laser Scanning microscope Zeiss LSM980 AiryScan 2 Axio Observer 7 SP with Definite Focus 3, driven by ZEN Blue v3.9, using Alpha Plan-APO 100 x objective and Axiocam 305 mono R2 camera.

Time-lapse imaging for FRAP experiments was performed using a Nikon Eclipse Ti-E microscope driven by MetaMorph Microscopy Automation and Image Analysis Software (Molecular Devices) and equipped with Yok-agawa CSU-X1-A1 confocal Spinning Disc unit and Prime BSI sCMOS camera (Teledyne Photometrics), using a Plan Apo VC 100x 1.4 NA objective. Targeted photobleaching was done using an ILas system (Roper Scientific France/PICT-IBiSA, Institute Curie).

All stacks were obtained at 0.25 µm intervals. Images were analyzed and processed using Fiji 1.54p and Imaris 10.2.0. For quantifications, the same laser power, exposure times and z-stack sizes were used within experiments. Time lapse stacks of developing embryos were acquired every 5 minutes for 4.75 hours. Images of PAC-1::GFP and LGL-1::GFP were processed using SAIBR for spectral auto fluorescence correction using the default settings (79).

### Fluorescence recovery after photobleaching (FRAP)

For FRAP assays, laser power was adjusted in each experiment to avoid complete photobleaching of the selected area. Photobleaching of epidermal DLG-1 was performed using a consistent laser setting for all animals and a photobleached area of ∼50 pixels in width. Time-lapse images were taken prior, during, and immediately after photobleaching. Recovery was followed at 1 s intervals for 200 s. Time-lapse movies were analyzed in Fiji. A 20 pixel wide line was drawn over the middle of the bleached area defined by the bleached region in the first post-bleach frame. For each time-lapse frame, the maximum intensity value within the bleached region was determined, and the background, defined by the mean intensity of a non-bleached region outside of the animal, was subtracted. The maximum intensities within the bleached region were corrected for acquisition photobleaching per frame using the background-subtracted maximum intensity of a similar non-bleached junction. FRAP recovery was calculated as the change in the corrected intensity values within the bleached region. The first frame after bleach was defined as 0% and the average maximum intensity of 8 frames before bleaching as 100%.

### DLG-1 peak measurement

Measurements of DLG-1 spread along the lateral membrane were done using a strain expressing PH::GFP to visualize the membranes and DLG-1::mCherry. To measure the length of the lateral membrane, a line was drawn over the PH::GFP signal and fluorescence intensities determined. The length of the membrane was defined as the length between 50% of the first and last peak heights. Next, the DLG-1 signal over the same line was measured. The DLG-1 signal length was defined as the area where fluorescence intensity was >40% of the difference between minimum and maximum signal intensities along the line.

### Apical curvature measurement

Apical curvature measurements were done using PH::GFP as a membrane marker. To measure the apical curvature of the epidermal cells, a segmented line was drawn over the apical domain of as many cells in a row as possible, defined as cells with clearly distinguishable apical and lateral domains (minimum 8 cells). Then a segmented line was drawn from junction to junction for the same cells, with straight lines between the junctions. The length of the apical domain line was divided by the length of the junction-to-junction line, giving one curvature value per embryo.

### Membrane length measurements

To measure the length of the apical, lateral and basal membranes of epidermal cells, segmented lines were drawn over the PH::GFP signal and their lengths and fluorescence intensities were determined. For apical and basal surfaces, lines were drawn along multiple cells in a row (minimum 8), and total lengths divided by the number of cells measured. The length of the lateral membranes were defined as the length between 50% of the first and last peak heights, measured individually and then averaged. The apical, lateral and basal measurements were taken from the same cells in one embryo.

### CDC-42 biosensor apical intensity measurements

To measure the CDC-42 biosensor intensity at the apical domain of the seam cells, a maximum intensity projection of a 3D stack covering the apical domain of the cells was generated. For cells with a clearly visible outline, a polygon selection tool was used to outline the area of the apical domain, as big as possible for every cell, that did not include the signal from the junctions. For each cell, the mean value was determined and background intensity from an area outside the embryo subtracted. The values were averaged for the whole embryo.

### Overall aPKC intensity measurements

To determine overall levels of aPKC::GFP in control and *pac-1(RNAi); lgl-1(mib201)* animals, an average intensity z-stack projection was made of the half of the embryo closest to the objective. A measurement area including the full embryo was drawn manually, and the average pixel intensity determined. The average intensity of a control area outside the embryo was subtracted to yield the final average aPKC intensity level.

### Statistical analysis

All statistical analysis was done using GraphPad Prism 10. For populations comparisons, a D’Agostino & Pearson test of normality was first performed to determine if the data was sampled from a Gaussian distribution. For data drawn from a Gaussian distribution, comparisons between two populations were done using an unpaired t test (with Welch’s correction if the standard deviations of the populations differed significantly), and for data that was not normally distributed, a Mann-Whitney test was performed. For multiple comparisons tests, for data drawn from Gaussian distribution, but with significantly different standard deviations, the Brown-Forsythe and Welch ANOVA test followed by Dunnett’s T3 multiple comparison test was used and for non-normally distributed data the Kruskal-Wallis test followed by Dunn’s multiple pairwise comparison test was performed.

## Supplemental material

This publication is accompanied by the following supplemental material:

- Figure S1. Genome-wide RNAi feeding *lgl-1* enhancer screen. Related to Figure 1.
- Figure S2. Junctional defects in *pac-1; lgl-1* embryos. Related to Figure 2.
- Figure S3. Loss of *lgl-1* and *pac-1* disrupts aPKC and LET-413 localization, but not CDC-42 activity. Related to Figure 3.
- Figure S4. Endogenously tagged PAC-1. Related to Figure 4.
- Video S1. Time-lapse videos of developing control and *pac-1(RNAi); lgl-1(mib201)* embryos. Related to Figure 1.
- Video S2. Localization of HMR-1 and DLG-1 in control and *pac-1(RNAi); lgl-1(mib201)* embryos. Related to Figure 2.
- Video S3. Time-lapse videos of two apical junction regions in a *pac-1(RNAi); lgl-1(mib201)* embryo. Related to Figure 2.
- Supplemental File S1. DNA sequence files of CRISPR alleles generated in this manuscript.
- Supplemental File S2. Raw data of all quantified figures.

## Data availability

All underlying numerical data for figures are provided in Supplemental File S2, including raw measurements, derived values, and the summary statistics plotted in the manuscript. All other data supporting the findings of this study are available within the paper and its supplementary information files. Raw microscopy image files and Adobe Illustrator source files with all embedded assets are publicly available at https://doi.org/10.24416/UU01-OK9BX0. *C. elegans* strains are available upon reasonable request.

## Acknowledgements

We thank members of the M. Boxem, S. van den Heuvel, S. Ruijtenberg, S. Suijkerbuijk and L. Braccioli groups for helpful discussions. We thank J. Nance, M. Labouesse, J. Rodriguez, N. Goehring, for strains and reagents. We thank the NYUAD Core Technology Platforms, in particular Nikolas Giakoumidis and Reza Rowshan, for assistance with the NYUAD high-throughput screening facility, and Nabil Rahiman for the RNAi screen scoring database. We thank H. Latonas for writing ImageJ analysis macros. We also thank Wormbase (83) and the Biology Imaging Center, Faculty of Sciences, Department of Biology, Utrecht University. Some strains were provided by the Caenorhabditis Genetics Center, which is funded by NIH Office of Research Infrastructure Programs (P40 OD010440).

## Author contributions

**Olga D. Jarosińska:** Conceptualization, Formal analysis, Investigation, Writing - Original Draft, Writing - Review & Editing, Visualization. **Amalia Riga:** Conceptualization, Formal analysis, Investigation, Writing - Review & Editing. **Hala Zahreddine Fahs:** Investigation, Supervision, Writing - Review & Editing. **Joren M. Woeltjes:** Investigation, Writing - Review & Editing. **Ruben Schmidt:** Investigation, Writing - Review & Editing. **Fathima S. Refai:** Investigation, Writing - Review & Editing. **Suma Gopinadhan:** Investigation, Writing - Review & Editing. **Kristin C. Gunsalus:** Supervision, Writing - Review & Editing, Funding acquisition. **Mike Boxem:** Conceptualization, Writing - Original Draft, Writing - Review & Editing, Supervision, Funding acquisition.

## References

1. Buckley C.E. and St Johnston D. Apical–basal polarity and the control of epithelial form and function. Nat Rev Mol Cell Biol. 23: 559–577 (2022). 10.1038/s41580-022-00465-y

2. Campanale J.P., Sun T.Y., and Montell D.J. Development and dynamics of cell polarity at a glance. J Cell Sci. 130: 1201–1207 (2017). 10.1242/jcs.188599

3. Rodriguez-Boulan E. and Macara I.G. Organization and execution of the epithelial polarity programme. Nat Rev Mol Cell Biol. 15: 225–242 (2014). 10.1038/nrm3775

4. St Johnston D. and Ahringer J. Cell polarity in eggs and epithelia: parallels and diversity. Cell. 141: 757–774 (2010). 10.1016/j.cell.2010.05.011

5. Betschinger J., Mechtler K., and Knoblich J.A. The Par complex directs asymmetric cell division by phosphorylating the cytoskeletal protein Lgl. Nature. 422: 326–330 (2003). 10.1038/na-ture01486

6. Hong Y. aPKC: the Kinase that Phosphorylates Cell Polarity. F1000Research. 7: F1000 Faculty Rev–903 (2018). 10.12688/f1000research.14427.1

7. Hurov J.B., Watkins J.L., and Piwnica-Worms H. Atypical PKC phosphorylates PAR-1 kinases to regulate localization and activity. Curr Biol CB. 14: 736–741 (2004). 10.1016/j.cub.2004.04.007

8. Yamanaka T., Horikoshi Y., Sugiyama Y., Ishiyama C., Suzuki A., Hirose T., Iwamatsu A., Shinohara A., and Ohno S. Mammalian Lgl Forms a Protein Complex with PAR-6 and aPKC Independently of PAR-3 to Regulate Epithelial Cell Polarity. Curr Biol. 13: 734–743 (2003). 10.1016/S0960-9822(03)00244-6

9. Yamanaka T., Horikoshi Y., Izumi N., Suzuki A., Mizuno K., and Ohno S. Lgl mediates apical domain disassembly by suppressing the PAR-3-aPKC-PAR-6 complex to orient apical membrane polarity. J Cell Sci. 119: 2107–2118 (2006). 10.1242/jcs.02938

10. Graybill C., Wee B., Atwood S.X., and Prehoda K.E. Partitioning-defective Protein 6 (Par-6) Activates Atypical Protein Kinase C (aPKC) by Pseudosubstrate Displacement*. J Biol Chem. 287: 21003–21011 (2012). 10.1074/jbc.M112.360495

11. Rodriguez J., Peglion F., Martin J., Hubatsch L., Reich J., Hirani N., Gubieda A.G., Roffey J., Fernandes A.R., St Johnston D., Ahringer J., and Goehring N.W. aPKC Cycles between Functionally Distinct PAR Protein Assemblies to Drive Cell Polarity. Dev Cell. 42: 400-415.e9 (2017). 10.1016/j.devcel.2017.07.007

12. Yamanaka T., Horikoshi Y., Suzuki A., Sugiyama Y., Kitamura K., Maniwa R., Nagai Y., Yamashita A., Hirose T., Ishikawa H., and Ohno S. PAR-6 regulates aPKC activity in a novel way and mediates cell-cell contact-induced formation of the epithelial junctional complex. Genes Cells. 6: 721–731 (2001). 10.1046/j.1365-2443.2001.00453.x

13. Achilleos A., Wehman A.M., and Nance J. PAR-3 mediates the initial clustering and apical localization of junction and polarity proteins during C. elegans intestinal epithelial cell polarization. Dev Camb Engl. 137: 1833–1842 (2010). 10.1242/dev.047647

14. Atwood S.X., Chabu C., Penkert R.R., Doe C.Q., and Prehoda K.E. Cdc42 acts downstream of Bazooka to regulate neuroblast polarity through Par-6/aPKC. J Cell Sci. 120: 3200–3206 (2007). 10.1242/jcs.014902

15. Dong W., Lu J., Zhang X., Wu Y., Lettieri K., Hammond G.R., and Hong Y. A polybasic domain in aPKC mediates Par6-dependent control of membrane targeting and kinase activity. J Cell Biol. 219: e201903031 (2020). 10.1083/jcb.201903031

16. Holly R.W., Jones K., and Prehoda K.E. A Conserved PDZ-Binding Motif in aPKC Interacts with Par-3 and Mediates Cortical Polarity. Curr Biol CB. 30: 893-898.e5 (2020). 10.1016/j.cub.2019.12.055

17. Lin D., Edwards A.S., Fawcett J.P., Mbamalu G., Scott J.D., and Pawson T. A mammalian PAR-3–PAR-6 complex implicated in Cdc42/Rac1 and aPKC signalling and cell polarity. Nat Cell Biol. 2: 540–547 (2000). 10.1038/35019582

18. McCaffrey L.M. and Macara I.G. The Par3/aPKC interaction is essential for end bud remodeling and progenitor differentiation during mammary gland morphogenesis. Genes Dev. 23: 1450–1460 (2009). 10.1101/gad.1795909

19. Martin-Belmonte F., Gassama A., Datta A., Yu W., Rescher U., Gerke V., and Mostov K. PTEN-Mediated Apical Segregation of Phosphoinositides Controls Epithelial Morphogenesis through Cdc42. Cell. 128: 383–397 (2007). 10.1016/j.cell.2006.11.051

20. Nunes de Almeida F., Walther R.F., Pressé M.T., Vlassaks E., and Pichaud F. Cdc42 defines apical identity and regulates epithelial morphogenesis by promoting apical recruitment of Par6-aPKC and Crumbs. Development. 146: dev175497 (2019). 10.1242/dev.175497

21. Peterson F.C., Penkert R.R., Volkman B.F., and Prehoda K.E. Cdc42 Regulates the Par-6 PDZ Domain through an Allosteric CRIB-PDZ Transition. Mol Cell. 13: 665–676 (2004). 10.1016/S1097-2765(04)00086-3

22. Qiu R.-G., Abo A., and Martin G.S. A human homolog of the C. elegans polarity determinant Par-6 links Rac and Cdc42 to PKCζ signaling and cell transformation. Curr Biol. 10: 697–707 (2000). 10.1016/S0960-9822(00)00535-2

23. Whitney D.S., Peterson F.C., Kittell A.W., Egner J.M., Prehoda K.E., and Volkman B.F. Binding of Crumbs to the Par-6 CRIB-PDZ Module Is Regulated by Cdc42. Biochemistry. 55: 1455–1461 (2016). 10.1021/acs.biochem.5b01342

24. Bilder D., Li M., and Perrimon N. Cooperative Regulation of Cell Polarity and Growth by Drosophila Tumor Suppressors. Science. 289: 113–116 (2000). 10.1126/science.289.5476.113

25. Hutterer A., Betschinger J., Petronczki M., and Knoblich J.A. Sequential Roles of Cdc42, Par-6, aPKC, and Lgl in the Establishment of Epithelial Polarity during Drosophila Embryogenesis. Dev Cell. 6: 845–854 (2004). 10.1016/j.devcel.2004.05.003

26. Tanentzapf G. and Tepass U. Interactions between the crumbs, lethal giant larvae and bazooka pathways in epithelial polarization. Nat Cell Biol. 5: 46–52 (2003). 10.1038/ncb896

27. Almagor L. and Weis W.I. Polarity protein Par6 facilitates the processive phosphorylation of Lgl via a dynamic interaction with aPKC. Commun Biol. 8: 967 (2025). 10.1038/s42003-025-08401-4

28. Earl C.P., Cobbaut M., Barros-Carvalho A., Ivanova M.E., Briggs D.C., Morais-de-Sá E., Parker P.J., and McDonald N.Q. Capture, mutual inhibition and release mechanism for aPKC–Par6 and its multisite polarity substrate Lgl. Nat Struct Mol Biol. 32: 729–739 (2025). 10.1038/s41594-024-01425-0

29. Herranz H., Stamataki E., Feiguin F., and Milán M. Self-refinement of Notch activity through the transmembrane protein Crumbs: modulation of γ-Secretase activity. EMBO Rep. 7: 297–302 (2006). 10.1038/sj.embor.7400617

30. Richardson E.C.N. and Pichaud F. Crumbs is required to achieve proper organ size control during Drosophila head development. Development. 137: 641–650 (2010). 10.1242/dev.041913

31. Tepaß U. and Knust E. Phenotypic and developmental analysis of mutations at thecrumbs locus, a gene required for the development of epithelia in Drosophila melanogaster. Rouxs Arch Dev Biol. 199: 189–206 (1990). 10.1007/bf01682078

32. Castiglioni V.G., Ramalho J.J., Kroll J., Stucchi R., Van Beuzekom H., Schmidt R., Altelaar M., and Boxem M. Characterization of the composition and functioning of the Crumbs complex in C. elegans. Developmental Biology; 2021. 10.1101/2021.08.10.455623

33. Waaijers S., Ramalho J.J., Koorman T., Kruse E., and Boxem M. The C. elegans Crumbs family contains a CRB3 homolog and is not essential for viability. Biol Open. 4: 276–284 (2015). 10.1242/bio.201410744

34. Chen J., Sayadian A.-C., Lowe N., Lovegrove H.E., and St Johnston D. An alternative mode of epithelial polarity in the Drosophila midgut. PLoS Biol. 16: e3000041 (2018). 10.1371/journal.pbio.3000041

35. Beatty A., Morton D., and Kemphues K. The C. elegans homolog of Drosophila Lethal giant larvae functions redundantly with PAR-2 to maintain polarity in the early embryo. Development. 137: 3995– 4004 (2010). 10.1242/dev.056028

36. Hoege C., Constantinescu A.-T., Schwager A., Goehring N.W., Kumar P., and Hyman A.A. LGL Can Partition the Cortex of One-Cell Caenorhabditis elegans Embryos into Two Domains. Curr Biol. 20: 1296– 1303 (2010). 10.1016/j.cub.2010.05.061

37. Boyd L., Guo S., Levitan D., Stinchcomb D.T., and Kemphues K.J. PAR-2 is asymmetrically distributed and promotes association of P granules and PAR-1 with the cortex in C. elegans embryos. Development. 122: 3075–3084 (1996). 10.1242/dev.122.10.3075

38. Montoyo-Rosario J.G., Armenti S.T., Zilberman Y., and Nance J. The Role of pkc-3 and Genetic Suppressors in Caenorhabditis elegans Epithelial Cell Junction Formation. Genetics. 214: 941–959 (2020). 10.1534/genetics.120.303085

39. Anderson D.C., Gill J.S., Cinalli R.M., and Nance J. Polarization of the C. elegans Embryo by RhoGAP-Mediated Exclusion of PAR-6 from Cell Contacts. Science. 320: 1771–1774 (2008). 10.1126/sci-ence.1156063

40. Klompstra D., Anderson D.C., Yeh J.Y., Zilberman Y., and Nance J. An instructive role for C. elegans E-cadherin in translating cell contact cues into cortical polarity. Nat Cell Biol. 17: 726–735 (2015). 10.1038/ncb3168

41. Zilberman Y., Abrams J., Anderson D.C., and Nance J. Cdc42 regulates junctional actin but not cell polarization in the Caenorhabditis elegans epidermis. J Cell Biol. 216: 3729–3744 (2017). 10.1083/jcb.201611061

42. Fic W., Bastock R., Raimondi F., Los E., Inoue Y., Gallop J.L., Russell R.B., and St Johnston D. RhoGAP19D inhibits Cdc42 laterally to control epithelial cell shape and prevent invasion. J Cell Biol. 220: e202009116 (2021). 10.1083/jcb.202009116

43. Cipriani P.G. and Piano F. RNAi Methods and Screening: RNAi Based High-Throughput Genetic Interaction Screening. Methods in Cell Biology. Elsevier; pp. 89–1112011. 10.1016/B978-0-12-544172-8.00004-9

44. Kamath R.S., Fraser A.G., Dong Y., Poulin G., Durbin R., Gotta M., Kanapin A., Le Bot N., Moreno S., Sohrmann M., Welchman D.P., Zipperlen P., and Ahringer J. Systematic functional analysis of the Caenorhabditis elegans genome using RNAi. Nature. 421: 231–237 (2003). 10.1038/na-ture01278

45. Chan E. and Nance J. Mechanisms of CDC-42 activation during contact-induced cell polarization. J Cell Sci. 126: 1692–1702 (2013). 10.1242/jcs.124594

46. Bossinger O., Klebes A., Segbert C., Theres C., and Knust E. Zonula Adherens Formation in Caenorhabditis elegans Requires dlg-1, the Homologue of the Drosophila Gene discs large. Dev Biol. 230: 29–42 (2001). 10.1006/dbio.2000.0113

47. Köppen M., Simske J.S., Sims P.A., Firestein B.L., Hall D.H., Radice A.D., Rongo C., and Hardin J.D. Cooperative regulation of AJM-1 controls junctional integrity in Caenorhabditis elegans epithelia. Nat Cell Biol. 3: 983–991 (2001). 10.1038/ncb1101-983

48. Shao X., Lucas B., Strauch J., and Hardin J. The adhesion modulation domain of Caenorhabditis elegans α-catenin regulates actin binding during morphogenesis. Fehon R, editor. Mol Biol Cell. 30: 2115–2123 (2019). 10.1091/mbc.E19-01-0018

49. Cox-Paulson E.A., Walck-Shannon E., Lynch A.M., Yamashiro S., Zaidel-Bar R., Eno C.C., Ono S., and Hardin J. Tropomodulin protects α-catenin-dependent junctional-actin networks under stress during epithelial morphogenesis. Curr Biol CB. 22: 1500–1505 (2012). 10.1016/j.cub.2012.06.025

50. Pasti G. and Labouesse M. Epithelial junctions, cytoskeleton, and polarity. WormBook. : 1–35 (2014). 10.1895/wormbook.1.56.2

51. Lynch A.M., Grana T., Cox-Paulson E., Couthier A., Cameron M., Chin-Sang I., Pettitt J., and Hardin J. A Genome-wide Functional Screen Shows MAGI-1 Is an L1CAM-Dependent Stabilizer of Apical Junctions in C. elegans. Curr Biol. 22: 1891–1899 (2012). 10.1016/j.cub.2012.08.024

52. Maiden S.L., Harrison N., Keegan J., Cain B., Lynch A.M., Pettitt J., and Hardin J. Specific Conserved C-terminal Amino Acids of Caenorhabditis elegans HMP-1/α-Catenin Modulate F-actin Binding Independently of Vinculin*. J Biol Chem. 288: 5694–5706 (2013). 10.1074/jbc.M112.438093

53. Pettitt J., Cox E.A., Broadbent I.D., Flett A., and Hardin J. The Caenorhabditis elegans p120 catenin homologue, JAC-1, modulates cadherin–catenin function during epidermal morphogenesis. J Cell Biol. 162: 15–22 (2003). 10.1083/jcb.200212136

54. Castiglioni V.G., Pires H.R., Rosas Bertolini R., Riga A., Kerver J., and Boxem M. Epidermal PAR-6 and PKC-3 are essential for larval development of C. elegans and organize non-centrosomal microtubules. Hobert O, Pfeffer SR, editors. eLife. 9: e62067 (2020). 10.7554/eLife.62067

55. Kagami M., Toh-e A., and Matsui Y. Sro7p, a Saccharomyces cerevisiae Counterpart of the Tumor Suppressor l(2)gl Protein, Is Related to Myosins in Function. Genetics. 149: 1717–1727 (1998). 10.1093/genetics/149.4.1717

56. Kumfer K.T., Cook S.J., Squirrell J.M., Eliceiri K.W., Peel N., O’Connell K.F., and White J.G. CGEF-1 and CHIN-1 Regulate CDC-42 Activity during Asymmetric Division in the Caenorhabditis elegans Embryo. Mol Biol Cell. 21: 266–277 (2010). 10.1091/mbc.E09-01-0060

57. Fievet B.T., Rodriguez J., Naganathan S., Lee C., Zeiser E., Ishidate T., Shirayama M., Grill S., and Ahringer J. Systematic genetic interaction screens uncover cell polarity regulators and functional redundancy. Nat Cell Biol. 15: 103–112 (2013). 10.1038/ncb2639

58. Jacob L., Opper M., Metzroth B., Phannavong B., and Mechler B.M. Structure of the I(2)gl gene of Drosophila and delimitation of its tumor suppressor domain. Cell. 50: 215–225 (1987). 10.1016/0092-8674(87)90217-0

59. Bulgakova N.A. and Knust E. The Crumbs complex: from epithelial-cell polarity to retinal degeneration. J Cell Sci. 122: 2587–2596 (2009). 10.1242/jcs.023648

60. Médina E., Lemmers C., Lane-Guermonprez L., and Le Bivic A. Role of the Crumbs complex in the regulation of junction formation in Drosophila and mammalian epithelial cells. Biol Cell. 94: 305–313 (2002). 10.1016/S0248-4900(02)00004-7

61. Segbert C., Johnson K., Theres C., van Fürden D., and Bossinger O. Molecular and functional analysis of apical junction formation in the gut epithelium of Caenorhabditis elegans. Dev Biol. 266: 17–26 (2004). 10.1016/j.ydbio.2003.10.019

62. Chalmers A.D., Pambos M., Mason J., Lang S., Wylie C., and Papalopulu N. aPKC, Crumbs3 and Lgl2 control apicobasal polarity in early vertebrate development. Development. 132: 977–986 (2005). 10.1242/dev.01645

63. Dodd M.E., Hatzold J., Mathias J.R., Walters K.B., Bennin D.A., Rhodes J., Kanki J.P., Look A.T., Hammerschmidt M., and Huttenlocher A. The ENTH domain protein Clint1 is required for epidermal homeostasis in zebrafish. Development. 136: 2591–2600 (2009). 10.1242/dev.038448

64. Klezovitch O., Fernandez T.E., Tapscott S.J., and Vasioukhin V. Loss of cell polarity causes severe brain dysplasia in Lgl1 knockout mice. Genes Dev. 18: 559–571 (2004). 10.1101/gad.1178004

65. Sonawane M., Martin-Maischein H., Schwarz H., and Nüsslein-Volhard C. Lgl2 and E-cadherin act antagonistically to regulate hemidesmosome formation during epidermal development in zebrafish. Development. 136: 1231–1240 (2009). 10.1242/dev.032508

66. Sripathy S., Lee M., and Vasioukhin V. Mammalian Llgl2 Is Necessary for Proper Branching Morphogenesis during Placental Development. Mol Cell Biol. 31: 2920–2933 (2011). 10.1128/MCB.05431-11

67. Clark B.S., Cui S., Miesfeld J.B., Klezovitch O., Vasioukhin V., and Link B.A. Loss of Llgl1 in retinal neuroepithelia reveals links between apical domain size, Notch activity and neurogenesis. Dev Camb Engl. 139: 1599–1610 (2012). 10.1242/dev.078097

68. Raman R., Damle I., Rote R., Banerjee S., Dingare C., and Sonawane M. aPKC regulates apical localization of Lgl to restrict elongation of microridges in developing zebrafish epidermis. Nat Commun. 7: 11643 (2016). 10.1038/ncomms11643

69. Sonawane M., Carpio Y., Geisler R., Schwarz H., Maischein H.-M., and Nuesslein-Volhard C. Zebrafish penner/lethal giant larvae 2 functions in hemidesmosome formation, maintenance of cellular morphology and growth regulation in the developing basal epidermis. Dev Camb Engl. 132: 3255–3265 (2005). 10.1242/dev.01904

70. Bii V.M., Rudoy D., Klezovitch O., and Vasioukhin V. Lethal giant larvae gene family (Llgl1 and Llgl2) functions as a tumor suppressor in mouse skin epidermis. BioRxiv Prepr Serv Biol. : 2023.03.06.531408 (2023). 10.1101/2023.03.06.531408

71. Stolpner N.J., Manzi N.I., Su T., and Dickinson D.J. Apical PAR protein caps orient the mitotic spindle in C. elegans early embryos. Curr Biol CB. 33: 4312-4329.e6 (2023). 10.1016/j.cub.2023.08.069

72. Chen W., Blanc J., and Lim L. Characterization of a promiscuous GTPase-activating protein that has a Bcr-related domain from Caenorhabditis elegans. J Biol Chem. 269: 820–823 (1994). 10.1016/S0021-9258(17)42184-3

73. Brenner S. The genetics of Caenorhabditis elegans. Genetics. 77: 71–94 (1974). 10.1093/ge-netics/77.1.71

74. Dickinson D.J., Pani A.M., Heppert J.K., Higgins C.D., and Goldstein B. Streamlined Genome Engineering with a Self-Excising Drug Selection Cassette. Genetics. 200: 1035–1049 (2015). 10.1534/genetics.115.178335

75. Ghanta K.S. and Mello C.C. Melting dsDNA Donor Molecules Greatly Improves Precision Genome Editing in Caenorhabditis elegans. Genetics. 216: 643–650 (2020). 10.1534/genetics.120.303564

76. Waaijers S., Muñoz J., Berends C., Ramalho J.J., Goerdayal S.S., Low T.Y., Zoumaro-Djayoon A.D., Hoffmann M., Koorman T., Tas R.P., Harterink M., Seelk S., Kerver J., Hoogenraad C.C., Bossinger O., Tursun B., van den Heuvel S., Heck A.J.R., and Boxem M. A tissue-specific protein purification approach in Caenorhabditis elegans identifies novel interaction partners of DLG-1/Discs large. BMC Biol. 14: 66 (2016). 10.1186/s12915-016-0286-x

77. Friedland A.E., Tzur Y.B., Esvelt K.M., Colaiácovo M.P., Church G.M., and Calarco J.A. Heritable genome editing in C. elegans via a CRISPR-Cas9 system. Nat Methods. 10: 741–743 (2013). 10.1038/nmeth.2532

78. Fahs H.Z., Refai F.S., Gopinadhan S., Moussa Y., Gan H.H., Hunashal Y., Battaglia G., Cipriani P.G., Ciancia C., Rahiman N., Kremb S., Xie X., Pearson Y.E., Butterfoss G.L., Maizels R.M., Esposito G., Page A.P., Gunsalus K.C., and Piano F. A new class of natural anthelmintics targeting lipid metabolism. Nat Commun. 16: 305 (2025). 10.1038/s41467-024-54965-w

79. Rodrigues N.T.L., Bland T., Borrego-Pinto J., Ng K., Hirani N., Gu Y., Foo S., and Goehring N.W. SAIBR: a simple, platform-independent method for spectral autofluorescence correction. Dev Camb Engl. 149: dev200545 (2022). 10.1242/dev.200545

80. Riga A., Cravo J., Schmidt R., Pires H.R., Castiglioni V.G., Van Den Heuvel S., and Boxem M. Caenorhabditis elegans LET-413 Scribble is essential in the epidermis for growth, viability, and directional outgrowth of epithelial seam cells. Nance J, editor. PLOS Genet. 17: e1009856 (2021). 10.1371/journal.pgen.1009856

81. Fielmich L.-E., Schmidt R., Dickinson D.J., Goldstein B., Akhmanova A., and Van Den Heuvel S. Optogenetic dissection of mitotic spindle positioning in vivo. eLife. 2018. 10.7554/eLife.38198

82. Kroll J.R., Remmelzwaal S., and Boxem M. CeLINC, a fluorescence-based protein-protein interaction assay in Caenorhabditis elegans. Genetics. 219: iyab163 (2021). 10.1093/genetics/iyab163

83. Sternberg P.W., Van Auken K., Wang Q., Wright A., Yook K., Zarowiecki M., Arnaboldi V., Becerra A., Brown S., Cain S., Chan J., Chen W.J., Cho J., Davis P., Diamantakis S., Dyer S., Grigoriadis D., Grove C.A., Harris T., et al. WormBase 2024: status and transitioning to Alliance infrastructure. Genetics. 227: iyae050 (2024). 10.1093/genetics/iyae050WW

