## Supplemental figures and legends for "LGL-1 and the RhoGAP protein PAC-1 redundantly polarize the *C. elegans* embryonic epidermis"

#### Contents

- Figure S1–S4
- Legends to supplemental videos S1–S3
- Legends to supplemental files S1 & S2
- Supplemental references

**Figure S1**

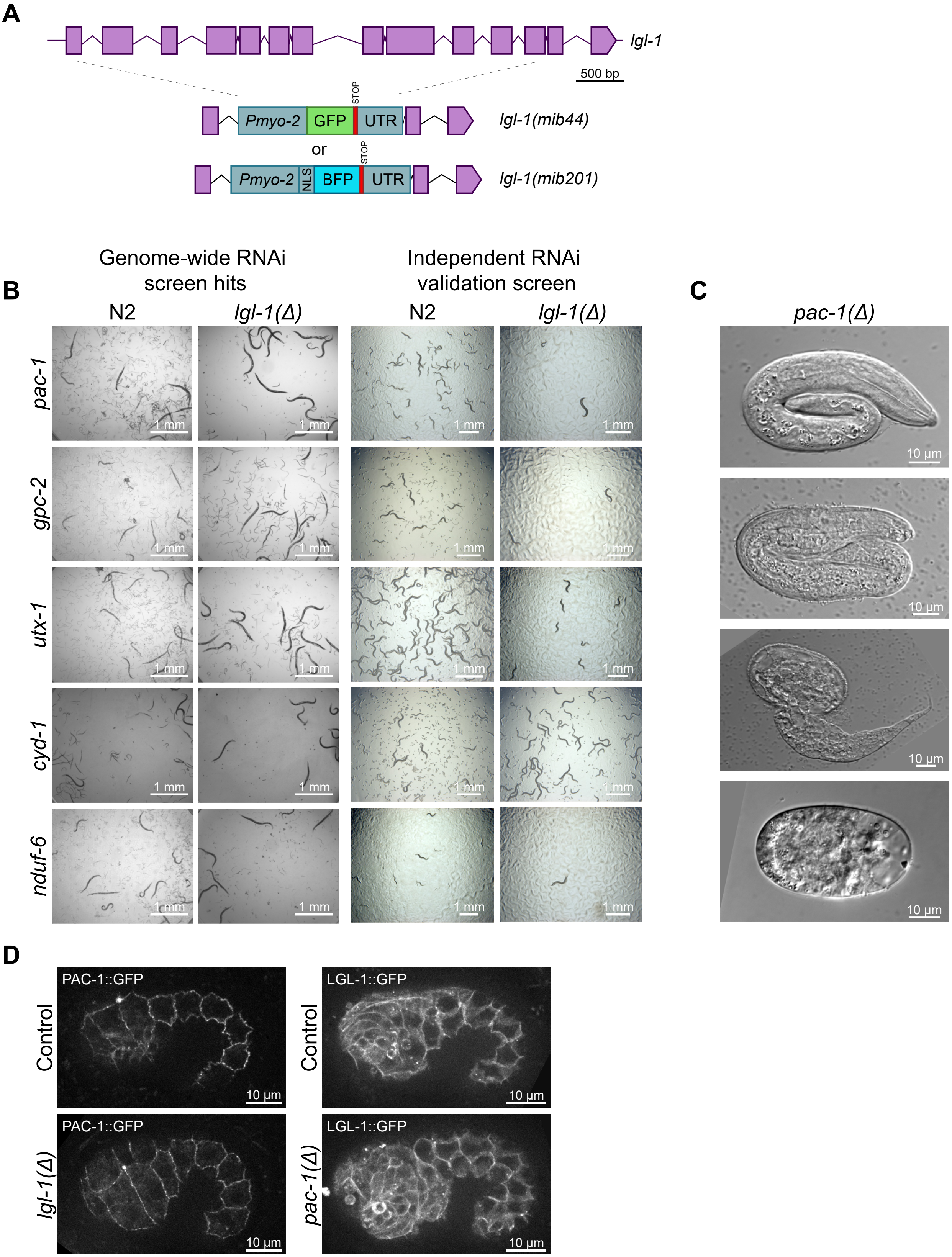


**Figure S1. Genome-wide RNAi feeding *lgl-1* enhancer screen.** (**A**) Genomic structure of *lgl-1* and the two *lgl-1* alleles – *lgl-1(mib44)* and *lgl-1(mib201)*. (**B**) Development of N2 and *lgl-1(mib44)* animals on feeding RNAi for the five confirmed *lgl-1(mib44)* enhancers *pac-1*, *gpc-2*, *utx-1*, *cyd-1*, and *nduf-6*. (**C**) Examples of terminal phenotypes of the 10% embryonic lethality caused by loss of *pac-1* alone. Strain used: BOX907. (**D**) Localization of PAC-1::GFP in *lgl-1(mib201)* and LGL-1::GFP in *pac-1(mib269)* embryos, as well as in controls. Images were processed using SAIBR for spectral auto fluorescence correction (Rodrigues et al., 2022). Strains used: NWG0285, BOX1102, BOX1075 and BOX1106.

**Figure S2**

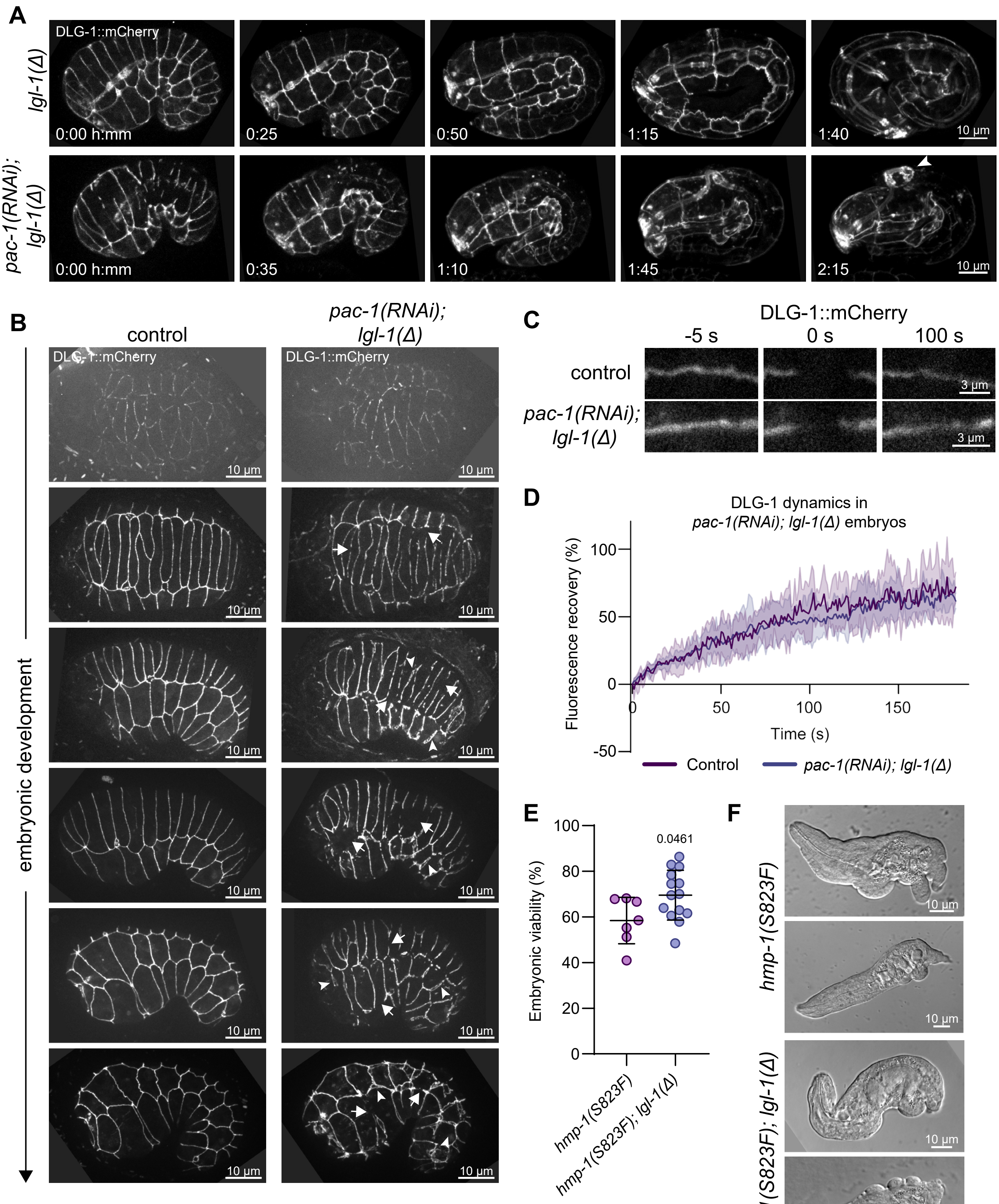


**Figure S2. Junctional defects in *pac-1; lgl-1* embryos.** (**A**) Stills from time-lapse videos showing the development of control and *pac-1(RNAi); lgl-1(mib201)* embryos. Embryos are visualized using an endogenously tagged DLG-1::mCherry fusion protein. Arrowhead indicates the site of epidermal rupture and the expulsion of the internal organs. Strain used: BOX899. (**B**) Images of DLG-1::mCherry in control and *pac-1(RNAi); lgl-1(mib201)* embryos taken at various stages of development. Strains used: BOX1076 and BOX1077. *Continued on next page.*

*Figure S2 legend continued.* (**C, D**) Stills from time-lapse imaging and FRAP curves of DLG-1::mCherry at the embryonic epidermis, in control and *lgl-1(mib201); pac-1(RNAi)* animals (n = 7 for control, n = 5 for *lgl-1(mib201); pac-1(RNAi)*). Thin lines and shading represent the mean ± SD. Strains used: BOX804, BOX940. (**E**) Embryonic viability of *hmp-1(S823F)* and *hmp-1(S823F); lgl-1(mib201)* mutant animals. Alleles *mib386* and *mib387* are identical S823F CRISPR mutations generated independently. Data is represented as mean ± SD and analyzed with a Brown-Forsythe and Welch ANOVA test followed by Dunnett's T3 multiple comparison test; p value shown on graph. Total plates counted in order of samples in graph: 7 and 14. Total progeny counted in order of samples in graph: 296 and 1066. (**F**) Examples of hatched embryos of indicated genotypes. While embryonic viable as judged by pharynx pumping, only a small fraction (~5%) of embryos survive to become fertile adults. Strains used: BOX1344 and BOX1348.

#### Figure S3

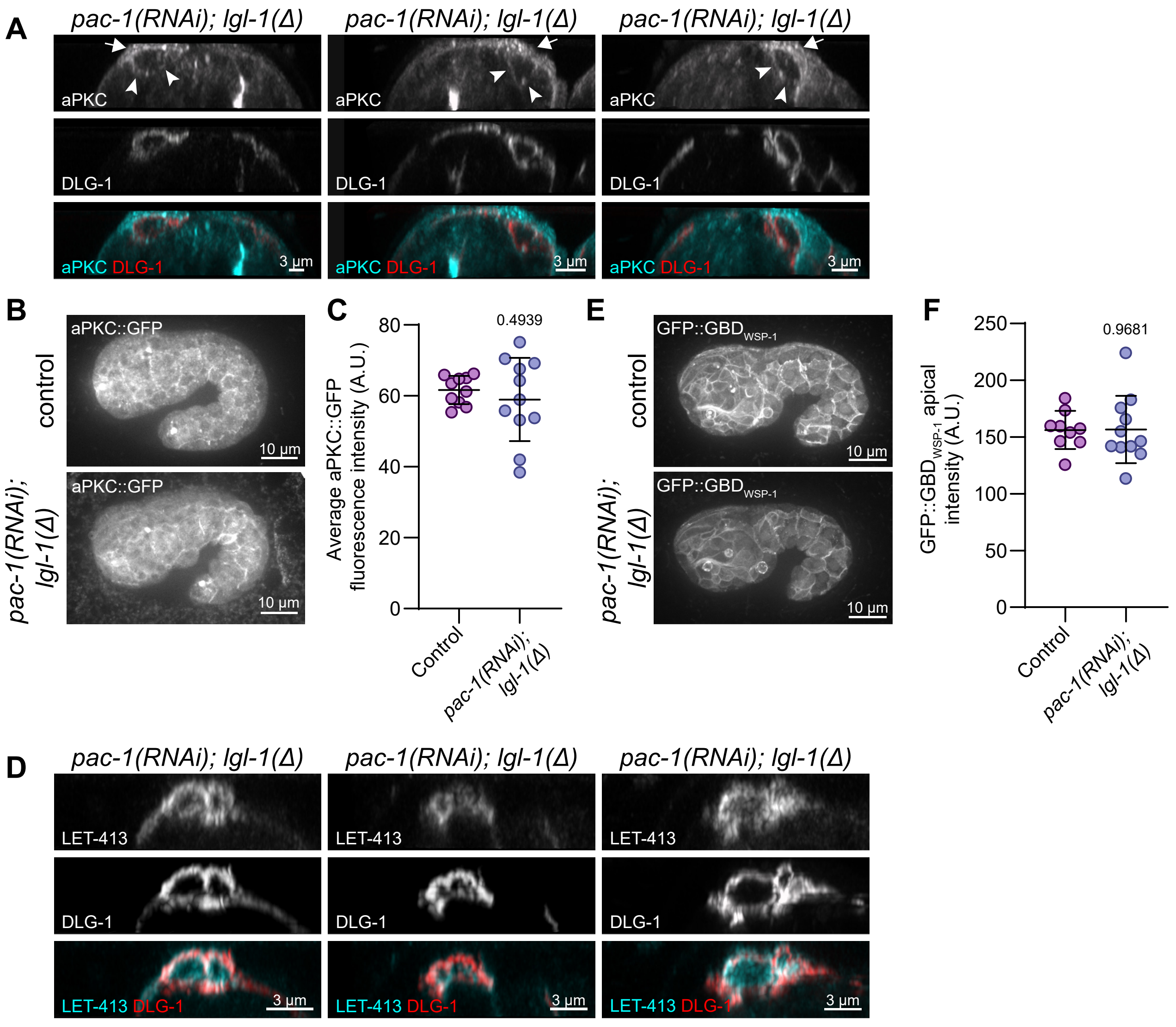


**Figure S3. Loss of *lgl-1* and *pac-1* disrupts aPKC and LET-413 localization, but not CDC-42 activity.** (**A**) Additional examples of mislocalization of aPKC and DLG-1 reporters in *pac-1; lgl-1* embryos. Strain used: BOX1161. (**B**) Average intensity projections of 3D stacks taken with a spinning disc confocal microscope showing the expression of aPKC::GFP in a control animal and in a *pac-1(RNAi); lgl-1(mib201)* animal. Strains used: BOX443 and BOX1045. (**C**) Quantification of total aPKC::GFP levels in images as in panel E. Data is represented as mean ± SD and analyzed using unpaired t test with Welch's correction; p value shown on graph. Total number of embryos quantified in order of samples in graph: 10 and 11. (**D**) Additional examples of mislocalization of LET-413 and DLG-1 reporters in *pac-1; lgl-1* embryos. Strain used: BOX1100. (**E**) Maximum intensity projections of 3D stacks taken with a spinning disc confocal microscope showing the localization of CDC-42 biosensor GFP::GBD$_{WSP-1}$. Strains used: FT1459 and BOX992. (**F**) Quantification of junctional GFP::GBD$_{WSP-1}$ localization in the embryonic epidermis of control and *pac-1; lgl-1* animals. Data is represented as mean ± SD and analyzed using unpaired t test with Welch's correction; p value shown on graph. Total number of embryos quantified in order of samples in graph: 8 and 11.

**Figure S4**

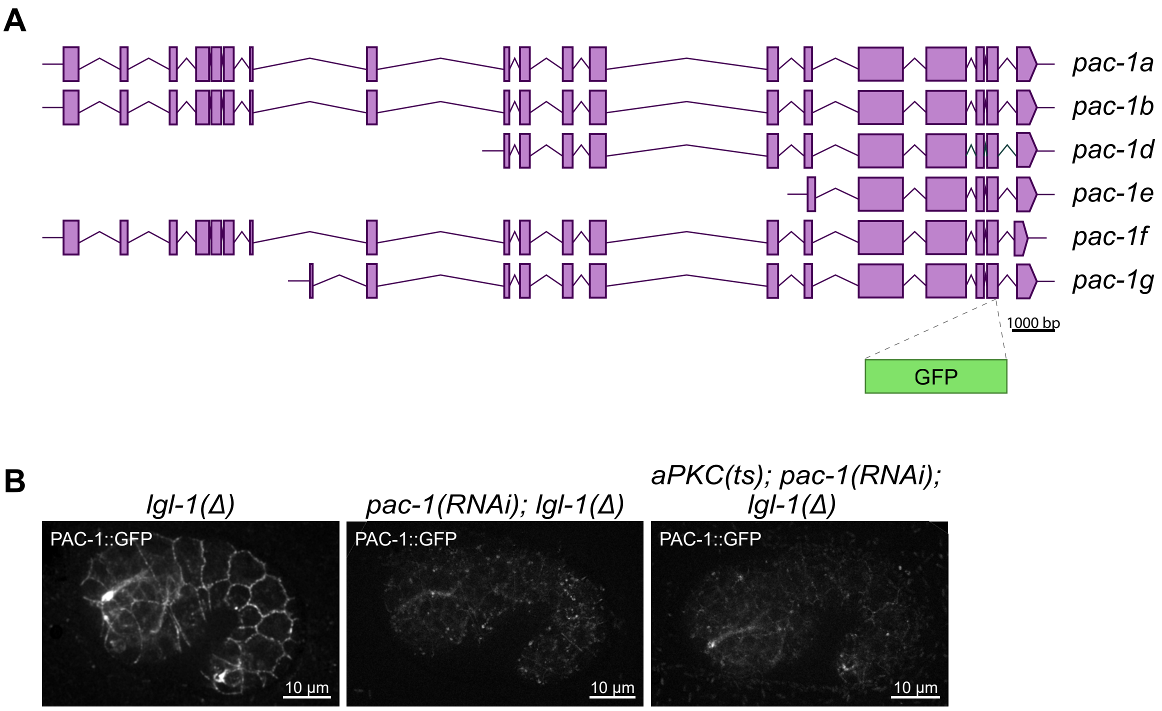


**Figure S4. Endogenously tagged PAC-1.** (**A**) Genomic structure of *pac-1* and the site of the GFP tag. (**B**) Maximum intensity projections of 3D stacks taken with spinning disc confocal microscope showing the localization of PAC-1::GFP. Strains used: BOX1177 and BOX1179. Images were processed using SAIBR for spectral auto fluorescence correction (Rodrigues et al., 2022).

### Supplemental legends

**Video S1. Time-lapse videos of developing control and *pac-1(RNAi); lgl-1(mib201)* embryos.** Membranes of the whole developing *C. elegans* embryo visualized using marker PH::GFP. Time lapse images were acquired every 5 minutes using Spinning Disc Fluorescent microscopy and the video is displayed at 6 frames/second. Arrowhead indicates site of epidermal rupture and expulsion of internal organs. Related to Figure 1.

**Video S2. Localization of HMR-1 and DLG-1 in control and *pac-1(RNAi); lgl-1(mib201)* embryos.** The epidermal junctions of control and *pac-1(RNAi); lgl-1(mib201)* embryos are visualized using DLG-1::mCherry (false-colored in magenta) and HMR-1::GFP (false-colored in green). The stacks were imaged using Airyscan confocal microscopy and the 3D fly-through renderings were generated in Imaris. Related to Figure 2.

**Video S3. Time-lapse videos of two apical junction regions in a *pac-1(RNAi); lgl-1(mib201)* embryo.** Both videos were taken in the same *pac-1(RNAi); lgl-1(mib201)* embryo. Left panel shows a epidermal junctional region with unaffected DLG-1::mCherry, while right panel shows a junctional region devoid of DLG-1::mCherry. The timelapse images were acquired using Spinning Disc microscopy, every 1 second and video is displayed as 6 frames/second. Related to Figure 2.

**Supplemental File S1. DNA sequence files of CRISPR alleles generated in this manuscript.** Each folder (AFD-1, DLG-1, LGL-1, and PAC-1) contains a final genomic sequence file and one or more '....design.dna' files which show primers and sgRNA sequences used annotated on the genomic sequence. Files are in SnapGene format and can be opened with the free viewer version of the software.

**Supplemental File S2. Raw data of all quantified figures.** Each sheet in the excel .xlsx file corresponds to the raw data for each of the quantifications found in all of the figures in the main and supplemental figures.

### Supplemental reference

**Rodrigues N.T.L., Bland T., Borrego-Pinto J., Ng K., Hirani N., Gu Y., Foo S., and Goehring N.W.** SAIBR: a simple, platform-independent method for spectral autofluorescence correction. *Dev Camb Engl.* **149**: dev200545 (2022). https://doi.org/10.1242/dev.200545
